# The transcription factor BCL11A restores differentiation potential to aged oligodendrocyte progenitor cells

**DOI:** 10.1101/2025.11.19.689239

**Authors:** Tanay Ghosh, Roey Baror, Chao Zhao, Amar Sharma, Wing Hei Au, András Lakatos, Nick Goldman, Robin JM Franklin

## Abstract

In young animals, oligodendrocyte progenitor cells (OPCs) undergo robust differentiation, progressing through stages to become pre-myelinating oligodendrocytes (Pre-OL) and ultimately myelinating oligodendrocytes (OLs). However, OPCs from aged animals have reduced differentiation ability. This disrupts myelin maintenance in the central nervous system (CNS), leading to a lack of remyelination following demyelinating injury and impaired adaptive myelination as a mechanism of learning. To uncover novel factors essential for restoring resilience in aged OPCs, we employed a data-driven approach involving the development of a computational pipeline, gSWITCH (accessible at: https://altoslabs.shinyapps.io/gSWITCH/), that allows for capture of precise dynamic gene expression patterns to pinpoint potential ‘switch’ genes during lineage progression. Using gSWITCH to identify potential switch genes crucial for OPCs, we conducted a comparative analysis of gene expression in OPCs isolated from young and aged animals. This revealed a group of transcription factors with decreased expression in aged OPCs. Further analysis of transcription factor binding site enrichment in OPC switches highlighted Bcl11a, a zinc finger transcription factor that could potentially serve as a master regulator of many switch genes. Ectopic overexpression of Bcl11a in aged OPCs did not enhance their proliferation; however, it significantly enhanced their differentiation into OLs. Overexpression of Bcl11a in aged mice, followed by demyelination injury in spinal cord white matter, significantly increased the differentiation of OPCs into OLs within the injury region compared to control aged mice. Furthermore, we found that Bcl11a is absent in invertebrates and has undergone pervasive purifying selection throughout vertebrate evolution, constraining its amino-acid sequence by eliminating deleterious mutations. Our study shows that reversing the age-related decline of this evolutionarily conserved factor in aged OPCs restores their impaired capacity for differentiation.

## Introduction

The regulation of gene expression during development is highly dynamic, guiding the transition from progenitor cells to terminally differentiated cell type (Peter and Davidson, 2011; Pope and Medzhitov, 2018). This temporal control ensures that appropriate gene expression programs are activated at each stage of lineage progression. Mapping these regulatory programs and identifying key driver within gene networks is essential for understanding how cell fate decisions are made and maintained. However, this developmental plasticity reduces with age, as progenitor cells progressively lose their ability to differentiate, impairing tissue homeostasis and repair (Rando et al., 2025; Brunet et al., 2023).

In the central nervous system (CNS), oligodendrocytes (OLs) are responsible for producing myelin, the lipid-rich sheath that insulates axons and accelerates electrical impulse conduction (Nave and Werner, 2014). In young animals, oligodendrocyte progenitor cells (OPCs) exhibit strong differentiation potential, transitioning through defined intermediate stages to generate mature, myelinating OLs. With ageing, however, this capacity declines: OPCs in older individuals show reduced ability to complete lineage progression, leading to compromised myelin maintenance, poor remyelination following demyelination injury and impaired adaptive myelination as a mechanism of learning (Franklin and ffrench-Constant, 2017). The molecular mechanisms underlying this age-related block in OPC differentiation remain poorly understood. Elucidating the regulatory factors that control OL lineage progression—and identifying their mis-regulation in aged OPCs—could uncover novel targets to restore resilience in these cells.

Here, we employed a data-driven approach to uncover novel factors essential for restoring resilience in aged OPCs. This involved the development of a computational pipeline, gSWITCH (accessible at: https://altoslabs.shinyapps.io/gSWITCH/), designed to capture precise dynamic gene expression patterns and identify potential ‘*switch*’ genes, including transcription factors, during lineage progression. Using this set of candidate switch genes, we performed a comparative analysis of gene expression between young and aged animals, which revealed a group of transcription factors downregulated in aged OPCs. Further analyses highlighted Bcl11a as a potential master regulator. Bcl11a expression was highest in OPCs and progressively decreased in Pre-OLs and OLs. Functional assays demonstrated that Bcl11a inhibition markedly reduced OPC differentiation into OLs. Overexpression of Bcl11a significantly enhanced differentiation in aged OPCs *in vitro* and following demyelination injury in the spinal cord of aged mice, without increasing OPC proliferation. Bcl11a is found only in vertebrates, and our molecular evolution analysis suggests that purifying selection has pervasively acted on this gene to preserve its amino acid sequences throughout vertebrate evolution. Our study showed that reversing the age-associated decline in expression of this evolutionarily conserved factor reinstates the differentiation capacity of aged OPCs.

## Results

### Identification of Switch genes during OL lineage progression

We performed RNA-seq analysis on OPC, Pre-OL and OL isolated from young rat (2-3 months) brains (**Figure 1A**). The OPC, Pre-OL, and OL samples were clearly distinguishable based on their global gene expression profiles (**Figure 1B**). The expression patterns of cell-type-specific markers were consistent with their being distinct OPC, Pre-OL, and OL populations (**Supplementary Figure 1,** see **Supplementary text** for detailed description).

**Figure 1.**
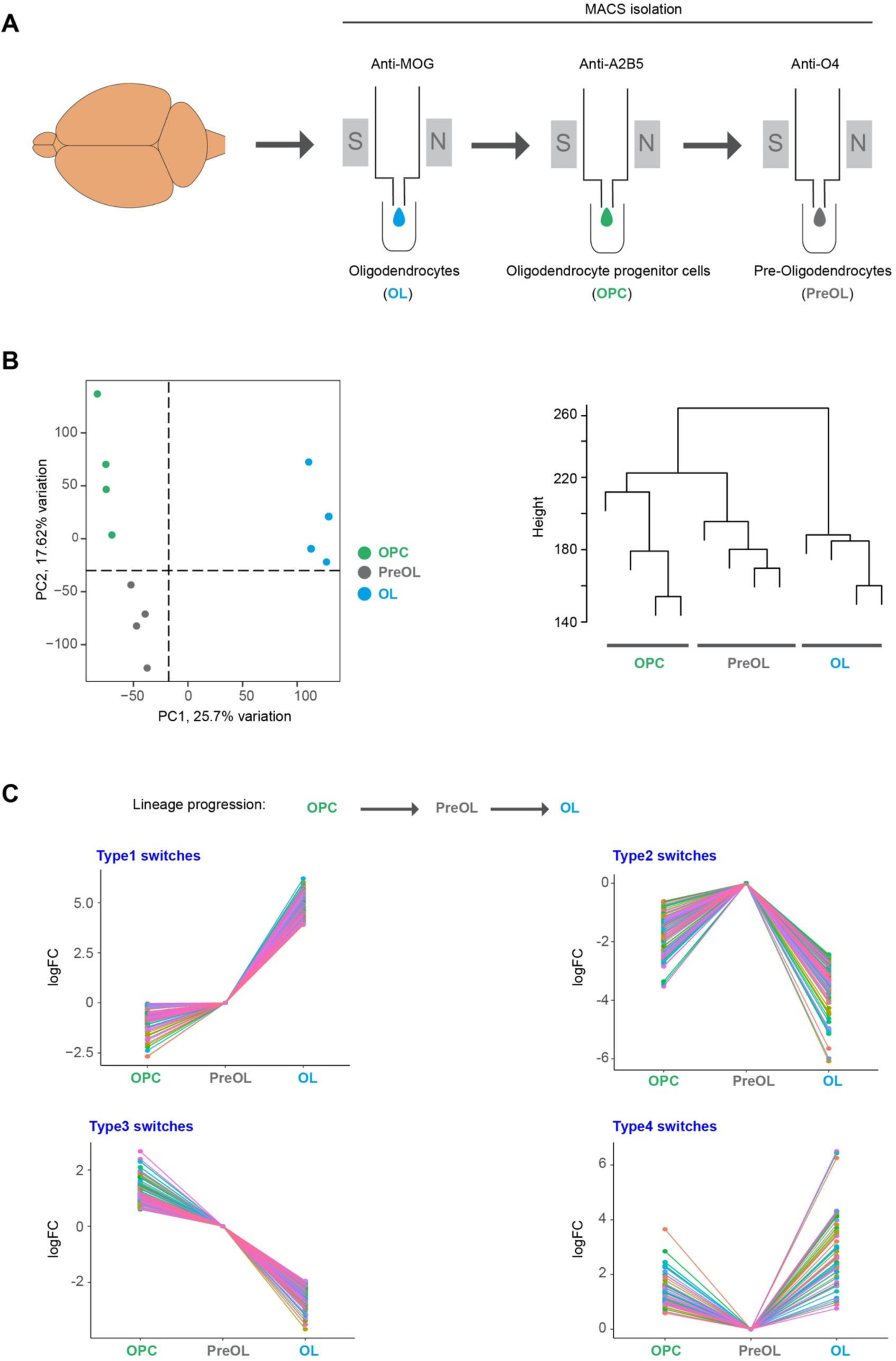
Dynamic regulation of genes during oligodendrocytes lineage progression. **A)** Schematic diagram showing magnetic-activated cell sorting (MACS)-based isolation of oligodendrocyte progenitor cells (OPCs), pre-oligodendrocytes (PreOLs), and oligodendrocytes (OLs) from young rat brains (2–3 months). The schema illustrates the order in which these cell types were most effectively isolated: mature OLs were isolated first using the OL-specific marker MOG, followed by OPCs using A2B5, and finally PreOLs using O4. **B)** Principal component analysis (PCA) (left) and average-linkage hierarchical clustering (right) based on global gene expression show that each cell type forms a distinct cluster. OPCs, PreOLs, and OLs were isolated from four different rats. **C)** Using gSWITCH, we identified four distinct patterns of switch genes during OL lineage progression. Each coloured line represents one gene and connects its log_2_fold-change values across the three oligodendrocyte lineage states: OPC, PreOL and OL. The connecting lines are used to visualize gene-wise patterns of expression change across these cell states. (**B-C**) GEO accession number of RNA-seq data: GSE301578. See also Supplementary Table 1, Supplementary Figure 1, Supplementary text, Supplementary Figure 3A.

To identify critical genes during oligodendrocyte lineage progression, we developed gSWITCH application (see Methods and **Supplementary Figure 2**).

Existing computational tools such as Monocle (Trapnell et al., 2014), tradeSeq (Van den Berge et al., 2020) and maSigPro (Nueda et al., 2014) are highly valuable for identifying genes with dynamic expression changes across pseudotime or time-course data. gSWITCH addresses a different question. It does not aim to infer trajectories. It works with a user-defined ordered series of biological states or time points and asks a more specific question — does this gene show a statistically supported “switch-like” change in expression as cells move through these states, and if so, what shape does that change take? It combines GLM-based statistical testing with criteria that capture where a gene reaches its highest or lowest expression and whether its expression changes steadily in one direction across the ordered series. To our knowledge, no existing tool combines significance testing with this type of explicit, shape-based classification into discrete, interpretable switch categories. gSWITCH sorts genes into four biologically meaningful patterns, rather than producing only a ranked list of significant genes based on pairwise comparisons between multiple conditions or states. gSWITCH also flags which of these switch genes are transcription factors, making it easier to prioritise candidates for follow-up experiments.

This biologist-friendly tool is freely available as a web application requiring no programming, works with experimental designs containing three or more stages or time points with at least two replicates per stage (no upper limit on either), and can be applied to bulk RNA-seq or to single-cell RNA-seq data aggregated as pseudobulk.

We identified four distinct dynamic patterns of gene expression, termed Type 1 to Type 4 switches (**Figure 1C**, **Supplementary Table 1, Supplementary Figure 3A**), during OL-lineage progression. These patterns are defined by the relative expression levels of specific gene sets across different cellular states. Disruptions to these expression levels can impair the coordinated dynamics and hinder lineage progression.

Among the four expression patterns, Types 3 and 4 are most characteristic of OPCs, as they show higher relative expression in OPCs than in the next lineage stage, PreOL. For Type 4, expression levels rise again from PreOL to OL, but note that they do not return to the levels observed in OPCs. Indeed, gSWITCH is designed to detect a statistically significant difference between the first and last cell states (here, OPC vs OL), based on a user-defined fold-change threshold (1.5 in our case) (**Supplementary Figure 2**).

### Altered expression of switches in aged OPCs

We next examined the regulation of type 3 and type 4 switches in aged OPCs. Our transcriptome meta-analysis showed clear differences between aged and young OPCs, identifying a set of genes that are consistently upregulated or downregulated in aged OPCs (**Figure 2A**, **Figure 2B**). A significant proportion of type 3 and type 4 switches were among these differentially expressed genes (135 out of 418 type 3 genes; 8 out of 45 type 4 genes) (**Figure 2C**, **Supplementary Figure 3B**). Notably, these switches were significantly downregulated in aged OPCs (**Figure 2C**), suggesting that altered regulation of these critical genes may impair the differentiation of aged OPCs into OLs.

**Figure 2.**
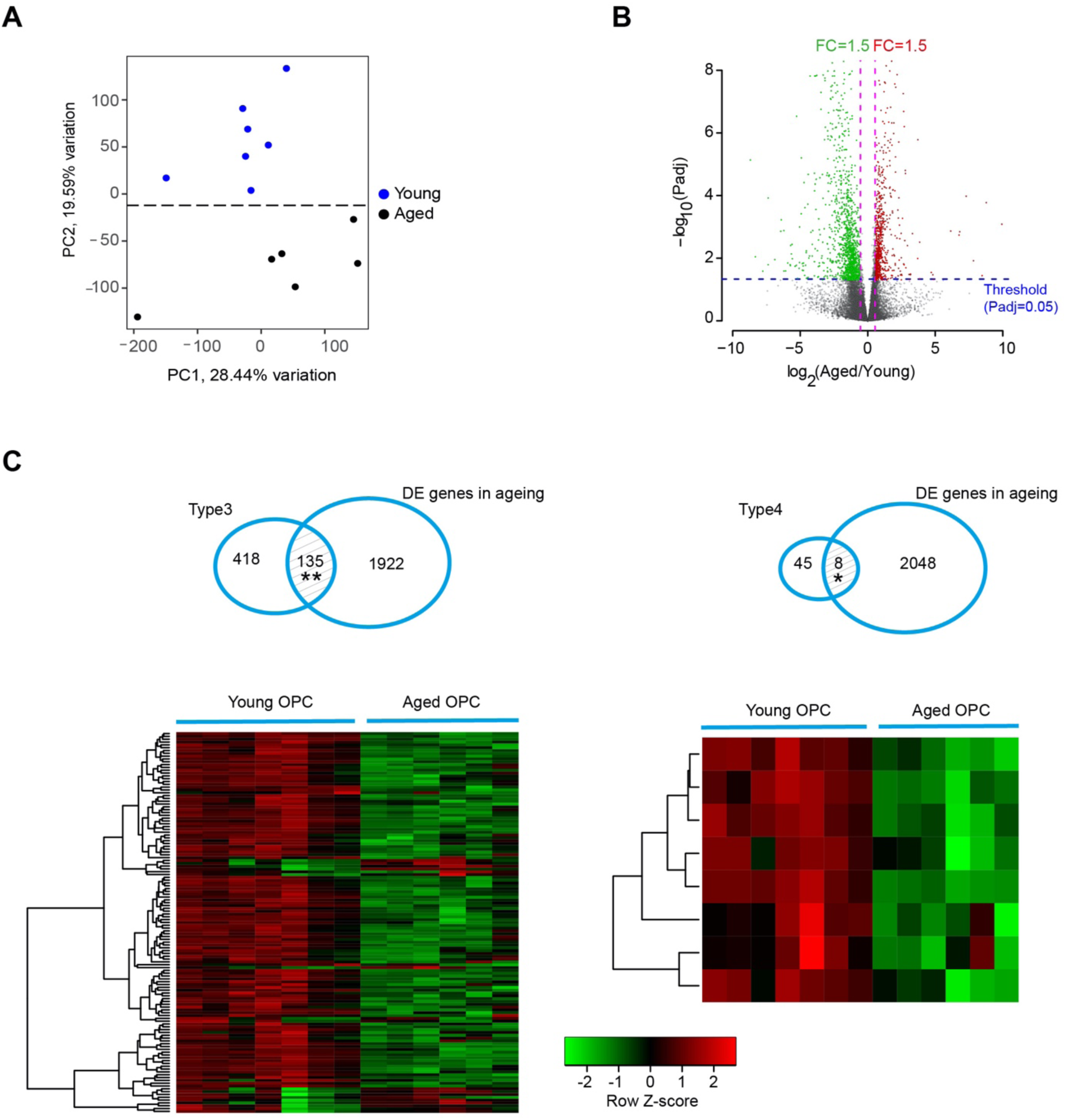
Down-regulation of switch genes due to ageing. **A)** Principal component analysis (PCA) shows that aged and young OPCs are distinguishable based on global gene expression (meta-analysis of RNA-seq datasets, GEO accession number: GSE303317). **B)** Volcano plot showing differentially expressed genes in aged OPCs compared to young OPCs. n= 6 (aged rats), 7 (young rats), FC: fold change; Padj: adjusted p-value (Benjamini–Hochberg FDR after Wald test). **C)** Venn diagram (top) showing the subset of Type 3 and Type 4 switch genes affected by ageing. Their expression patterns are shown in the heat map (bottom), where we observed down-regulation with ageing. * P < 0.05, ** P < 0.001, Chi-square test (two sided) with Yates’ correction. DE: Differentially expressed. DE gene list is accessible through GEO accession: GSE303317. See also Supplementary Table 1, Supplementary Figure 3B.

### Bcl11a is identified as a potential master switch

The orchestrated action of transcription factors (TFs) gradually drives the developmental program (Spitz F and Furlong EE, 2012). Using gSWITCH, we identified switches that are TFs and found 67 TFs among type 3 and type 4 switches (**Supplementary Table 2**).

Because a single transcription factor (TF) can regulate the expression of many genes, we aimed to identify TFs that may regulate type 3 and type 4 switches. TF binding site enrichment analysis using two chromatin immunoprecipitation (ChIP)-sequencing experimental evidence-based databases, ENCODE and ReMap2, predicted 27 TFs common to both datasets (**Figure 3A, Supplementary Table 3**). We then examined whether any of these 27 TFs were present in type 3 and type 4 switches and identified five [Zinc finger protein B-cell CLL/Lymphoma 11A (Bcl11a) (also known as CTIP1 (COUP-TF–interacting protein 1), CCAAT/enhancer-binding protein beta (Cebpb), Forkhead box M1 (Foxm1), Jun Proto-Oncogene, AP-1 Transcription Factor Subunit (Jun), and PR Domain Zinc Finger Protein 1 (Prdm1)], all of which were defined as type 3 switches (**Figure 3A**, **Figure 3B, Supplementary Table 3**). Among these, only Bcl11a and Foxm1 were significantly downregulated in aged OPCs (**Figure 3B**). We focused further on Bcl11a, as its magnitude of change being greater than that of Foxm1 (**Figure 3B**).

**Figure 3.**
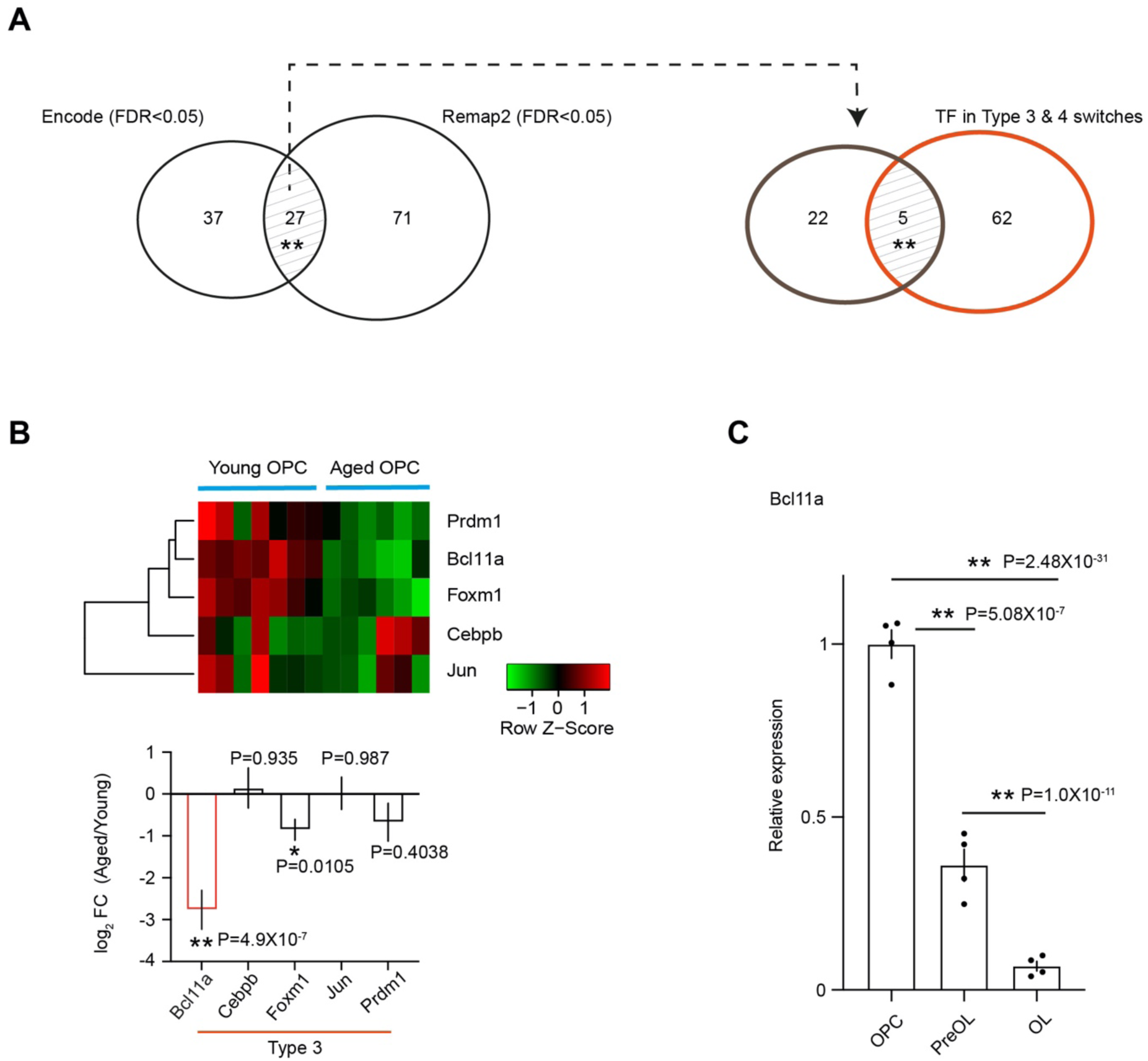
Transcription factor Bcl11a-mediated network potentially affected duuring ageing. **A)** Transcription factor binding site enrichment analysis using Type 3 and Type 4 switch genes as input identified 27 transcription factors (TFs) detected in both ENCODE and ReMap2, which are ChIP-seq–based experimental databases. Among these 27 TFs, we found 5 TFs that are themselves present as Type 3 or Type 4 switches (TF switches extracted from gSWITCH). ** P < 0.001, hypergeometric test. **B)** Heat map (top) showing expression (RNA-seq, GEO accession number: GSE303317) of these 5 TFs. Their differential expression in aged OPCs relative to young OPCs plotted as log_2_FC (bottom). FC: Fold change, n= 6 (aged rats), 7 (young rats), mean±SEM, * adjusted p-value < 0.05, ** adjusted p-value < 1×10⁻⁶, Benjamini–Hochberg FDR after Wald test. Note: All five TFs are classified as Type 3 switches, and Bcl11a shows the strongest repression in aged OPCs, with the highest statistical significance. **C)** Bcl11a, as a Type 3 TF switch, shows significantly repressed expression as OL lineage progresses. DESeq2-normalised values relative to OPCs plotted. n= 4 different rats, mean±SEM, ** adjusted p-value < 1×10⁻⁶, Benjamini–Hochberg FDR after Wald test. GEO accession number of RNA-seq data: GSE301578. See also Supplementary Table 2, Supplementary Table 3, Supplementary Figure 4.

The expression of Bcl11a followed a typical type 3 pattern: it was highest in OPCs and gradually decreased during differentiation, reaching significantly lower levels in OLs (**Figure 3C, Supplementary Figure 4A**). We therefore experimentally tested whether Bcl11a plays a functional role in OPC-to-OL differentiation, and whether restoring its expression can rescue the impaired differentiation capacity of aged OPCs.

### Inhibition of Bcl11a in OPCs markedly reduced their differentiation

Since Bcl11a expression decreases as OPCs differentiate into OLs (**Figure 3C, Supplementary Figure 4A**), we addressed the hypothesis that, if Bcl11a acts as a master switch for lineage progression, its early expression in OPCs would be functionally required for differentiation to proceed. To test this, we used small interfering RNA (siRNA)-mediated knockdown. Transfection with siRNA targeting Bcl11a (siBcl11a) reduced its expression by approximately 79% in proliferating OPCs (**Figure 4A**). Using the 5-ethynyl-2’-deoxyuridine (EdU) incorporation assay, we found no significant effect of Bcl11a knockdown on OPCs proliferation (**Figure 4B**). However, Bcl11a inhibition led to a substantial (∼77%) reduction in differentiation, as measured by immunostaining for myelin basic protein (MBP), a marker of OL differentiation (**Figure 4C**). These findings suggest that as a type 3 switch Bcl11a function is essential for priming OPCs for efficient differentiation.

**Figure 4.**
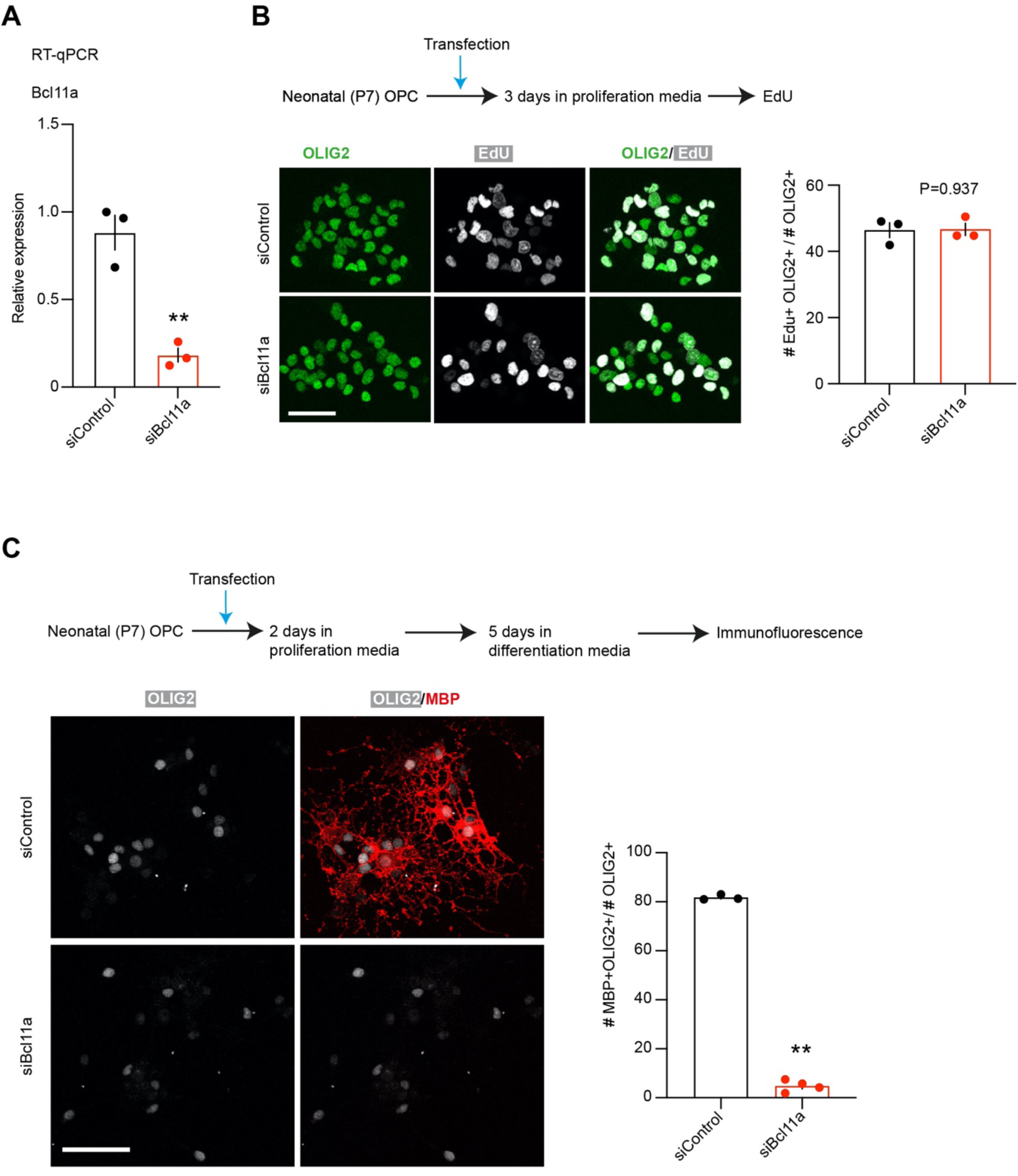
Effect of Bcl11a inhibition on OPCs proliferation and differentiation. **A)** Reverse transcription (RT)–qPCR analysis of Bcl11a expression in OPCs transfected with siRNA targeting Bcl11a. siBcl11a: siGENOME Rat Bcl11a; siControl: siGENOME non-targeting control siRNA (Dharmacon™). **B) Effect on proliferation.** Left: Immunofluorescence analysis using antibodies to OLIG2 and EdU detection (Click-iT protocol), followed by EdU proliferation assay. Right: Quantification of EdU⁺OLIG2⁺ cells plotted as a percentage of OLIG2⁺ cells. P: p-value, Student’s t test (unpaired and two tailed) **C) Effect on differentiation.** Left: Immunofluorescence analysis using antibody to MBP, OLIG2 upon inhibition of Bcl11a in OPCs. Right: quantification of MBP+ cells were plotted as a percentage of OLIG2+ cells. **(A-C)** n = 3 independent experiments (each time 4 P7 brains were pooled for OPC isolation). mean±SEM. ** p<0.01, Student’s t test (unpaired, two tailed); The symbol “#” indicates cell count. **(B-C)** scale bar: 22 µm.

### Transient overexpression of Bcl11a in aged OPCs improves differentiation

As noted, our RNA-seq analysis revealed reduced Bcl11a expression in aged OPCs (**Figure 3B, Supplementary Figure 4B**). To investigate whether restoring Bcl11a levels could overcome their impaired differentiation capacity, we expressed Bcl11a in aged OPCs (isolated from 20-24-month-old rats). We first performed *in vitro* transcription to generate Bcl11a mRNA (mBcl11a) and confirmed the fidelity of its protein expression by western blot analysis (**Supplementary Figure 5A**). mBcl11a was subsequently transfected into aged OPCs; Gfp mRNA (mGfp) served as a control. Transfection efficiency was approximately 77% (**Supplementary Figure 5B**). Similar to Bcl11a knockdown, mBcl11a expression had no significant effect on aged OPC proliferation (**Figure 5A**). However, culturing these OPCs in differentiating media led to an approximately 54% increase in OL numbers (**Figure 5B**), indicating that Bcl11a expression in OPCs is critical for their progression toward the oligodendrocyte lineage.

**Figure 5.**
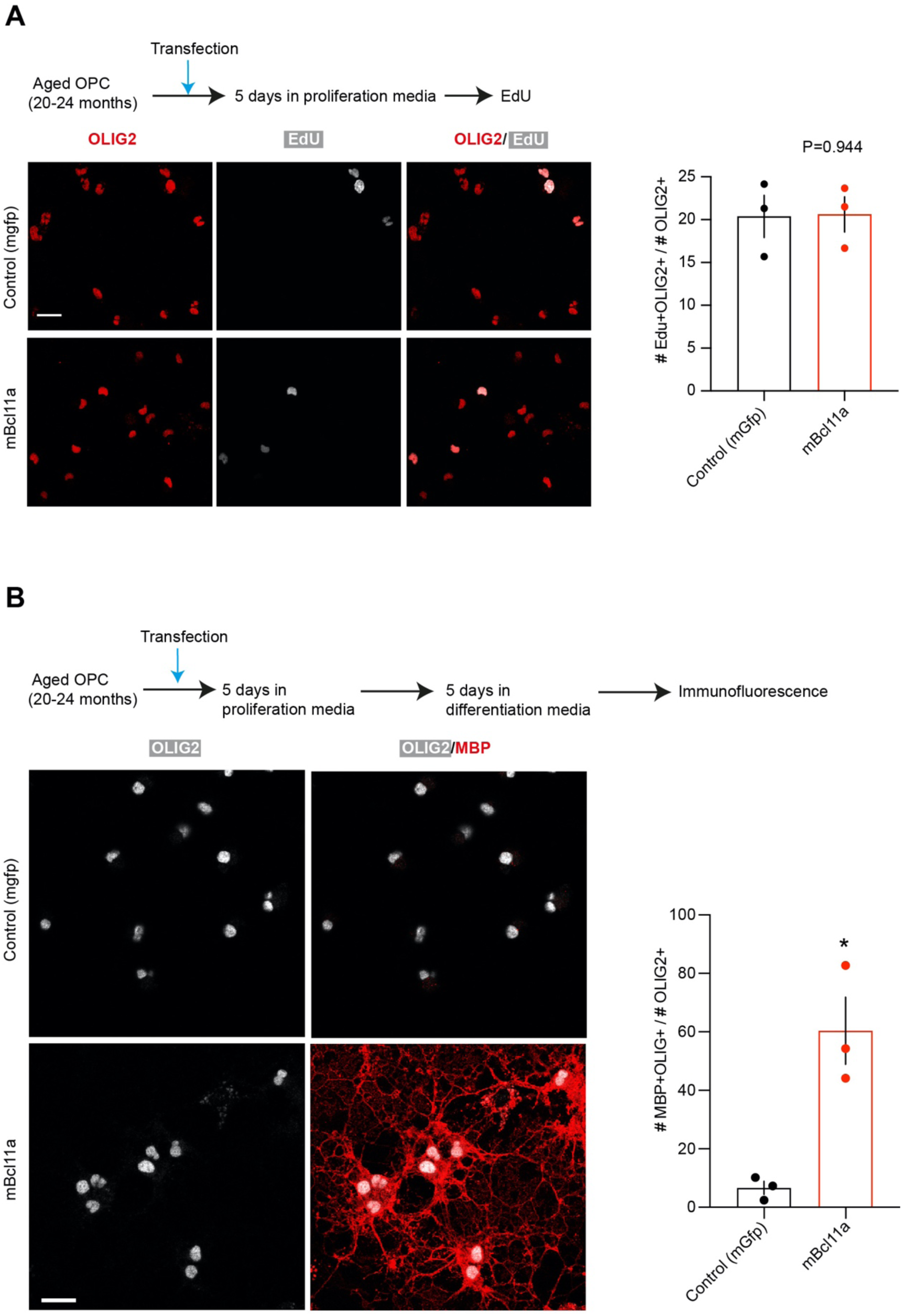
Effect of Bcl11a transient over-expression on proliferation and differentiation of aged OPCs. A) Effect on proliferation. Left: Immunofluorescence analysis using antibodies to OLIG2 and EdU detection (Click-iT protocol), followed by EdU proliferation assay. Right: Quantification of EdU⁺OLIG2⁺ cells plotted as a percentage of OLIG2⁺ cells. P: p-value, Student’s t test (unpaired, two tailed). **B) Effect on differentiation.** Left: Immunofluorescence analysis using antibodies to MBP and OLIG2 after transfection of Bcl11a mRNA (mBcl11a) or Gfp mRNA (mGfp; control) in aged OPCs. Right: Quantification of MBP⁺ cells plotted as a percentage of OLIG2⁺ cells. * P<0.05, Student’s t test (unpaired, two tailed). **(A-B)** n = 3 independent experiments (each using OPCs isolated from pooled brains of 2 aged rats). mean ± SEM. The symbol “#” indicates cell count. scale bar: 22 µm.

We next tested the effect of Bcl11a expression in OPCs in aged mice (18-month-old) using a model of focal demyelination in spinal cord white matter. We injected an adeno-associated virus (AAV) carrying a Sox10 driven Bcl11a-P2A-gfp construct into the tail vein of 18-month-old mice (**Figure 6A**). This design ensures Sox10-driven expression of both Bcl11a and Gfp via a self-cleaving P2A peptide. As a control, we injected AAV carrying a Sox10-driven Gfp construct. One month after AAV injection, we induced spinal cord lesions using lysolecithin. Fourteen days later, the spinal cords were harvested. We found that nearly all GFP immune-positive cells were also OLIG2 positive (98.6% in controls, total GFP+ 4’,6-diamidino-2-phenylindole [DAPI]+ cells counted: 193; 98% in Bcl11a, total GFP+ DAPI+ cells counted: 163), indicating strong oligodendrocyte lineage fidelity (**Supplementary Figure 5C**). Our protocol infected approximately 45% of GFP+OLIG2+ cells (190 GFP+OLIG2+ out of 422 OLIG2+ cells counted in control; 160 GFP+ OLIG2+ out of 357 OLIG2+ cells counted in Bcl11a) (**Supplementary Figure 5C**). We performed RNA *in situ* hybridization using a probe specific for proteolipid protein 1 (Plp1), a marker of late oligodendrocyte differentiation. We observed no difference in total OLIG2+ cells in the lesion area but a significant increased number of OLIG2+ cells that were Plp1+ in Bcl11a samples (**Supplementary Figure 6**; **Figure 6B**), suggesting Bcll1a restores the differentiation capacity of aged OPCs recruited to the injury area.

**Figure 6.**
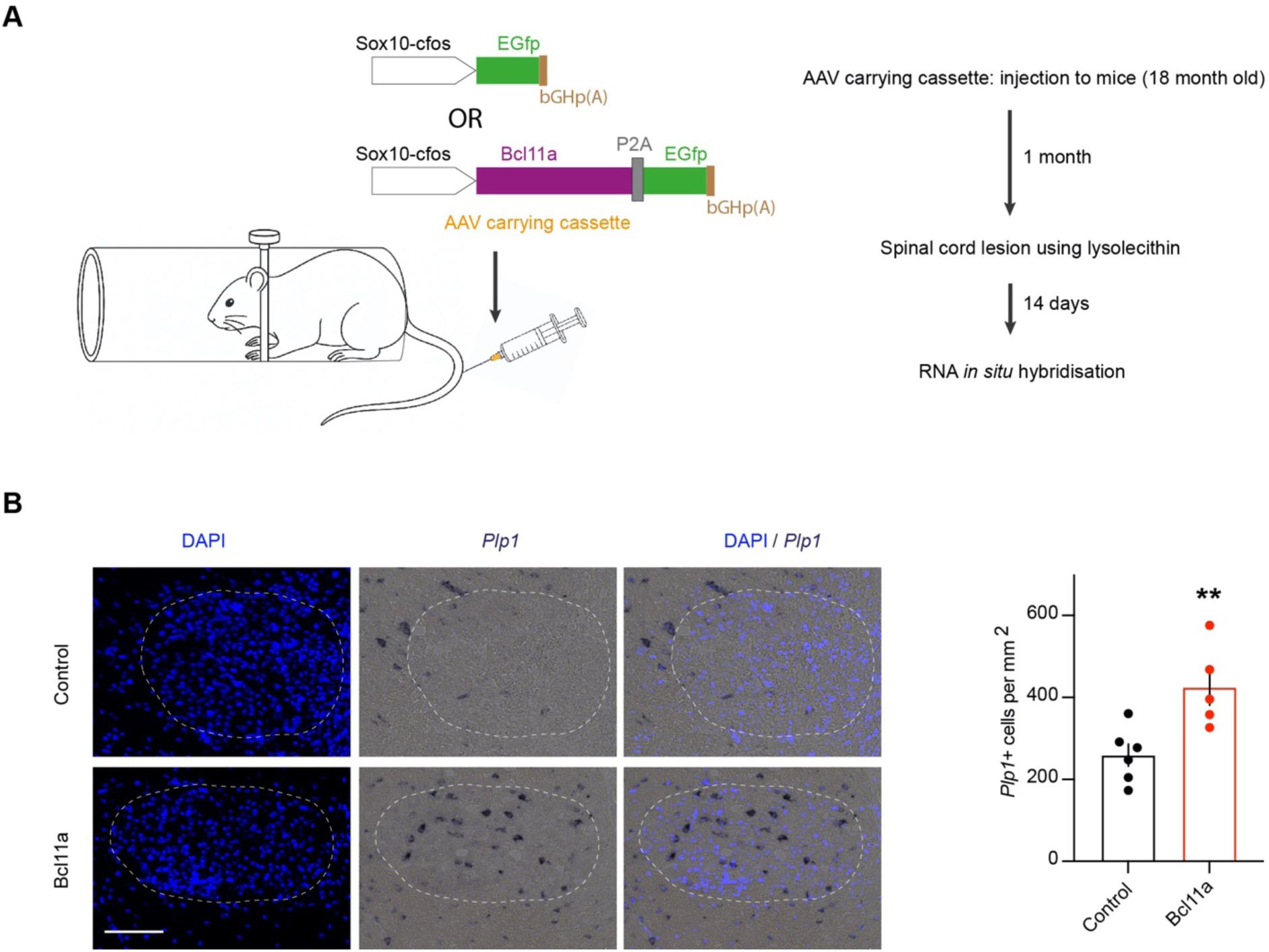
OL lineage specific Bcl11a expression in aged mice. **A)** Left: SOX10-driven construct carrying either Gfp alone (control) or Bcl11a linked to Gfp via a self-cleaving peptide element (P2A). This construct ensures SOX10-driven expression of both Bcl11a and Gfp. P2A: Porcine teschovirus-1 2A peptide, bGH p(A): bovine growth hormone poly(A) signal. The AAV containing this construct was delivered to 18-month-old mice via tail-vein injection. Right: schematic showing the experimental timeline. **B)** RNA *in situ* analysis of *Plp1*. In the lesioned area, the number of *Plp1*⁺ cells per mm² was counted and plotted. n=5-6 mice, mean±SEM, ** P<0.01, Student’s t test (unpaired, two tailed). scale bar: 100 µm. The white dotted line demarcates the lesion area.

### Bcl11a has undergone purifying selection during vertebrate evolution

Given that Bcl11a functions as an important regulatory transcription factor switch during oligodendrocyte (OL) lineage progression and that its expression level is associated with functional vulnerability in ageing, we used computational molecular evolution approaches to investigate whether selective constraints during evolution have acted on its codon sequence. We included twenty-three vertebrate species, including humans (**Supplementary Table 4**). We aimed to determine whether such constraints have preserved the amino acid sequence by eliminating deleterious mutations through purifying selection. Bcl11a is absent in invertebrate genomes (UCSC Genome Browser search), but orthologs are present across all examined vertebrate species. The ratio of nonsynonymous (dN) to synonymous (dS) substitution rates — known as the dN/dS ratio — is a widely used metric in molecular evolution to evaluate the selective pressures on protein-coding genes (Yang Z, 2007, Anisimova M and Kosiol C, 2009). As shown in **Table 1**, our analysis using multiple evolutionary models consistently yielded dN/dS values below 1, providing strong evidence of pervasive purifying selection acting on Bcl11a across the vertebrate lineage.

**Table 1:**
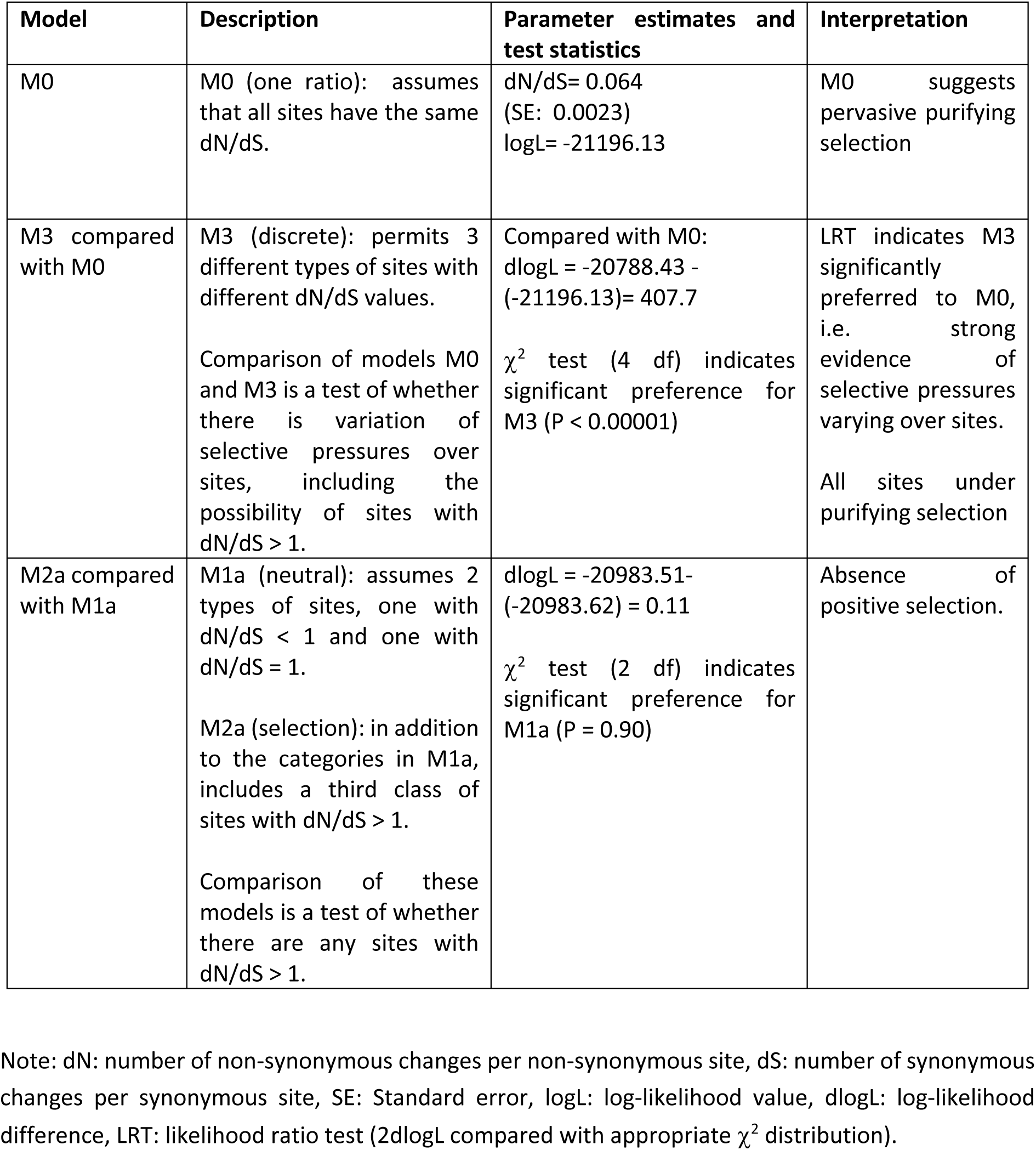
Test for evolutionary selection on Bcl11a.

| Model | Description | Parameter estimates and test statistics | Interpretation |
| --- | --- | --- | --- |
| M0 | M0 (one ratio): assumes that all sites have the same dN/dS. | dN/dS= 0.064<br>(SE: 0.0023)<br>logL= -21196.13 | M0 suggests pervasive purifying selection |
| M3 compared with M0 | M3 (discrete): permits 3 different types of sites with different dN/dS values.<br>Comparison of models M0 and M3 is a test of whether there is variation of selective pressures over sites, including the possibility of sites with dN/dS > 1. | Compared with M0:<br>dlogL = -20788.43 - (-21196.13)= 407.7<br>$\chi^2$ test (4 df) indicates significant preference for M3 (P < 0.00001) | LRT indicates M3 significantly preferred to M0, i.e. strong evidence of selective pressures varying over sites.<br>All sites under purifying selection |
| M2a compared with M1a | M1a (neutral): assumes 2 types of sites, one with dN/dS < 1 and one with dN/dS = 1.<br>M2a (selection): in addition to the categories in M1a, includes a third class of sites with dN/dS > 1.<br>Comparison of these models is a test of whether there are any sites with dN/dS > 1. | dlogL = -20983.51- (-20983.62) = 0.11<br>$\chi^2$ test (2 df) indicates significant preference for M1a (P = 0.90) | Absence of positive selection. |

Note: dN: number of non-synonymous changes per non-synonymous site, dS: number of synonymous changes per synonymous site, SE: Standard error, logL: log-likelihood value, dlogL: log-likelihood difference, LRT: likelihood ratio test (2dlogL compared with appropriate $\chi^2$ distribution).

## Discussion

Dynamically regulated genes during transitions from one cellular stage to another offer important information about the regulatory mechanisms governing differentiation (Spitz and Furlong, 2012; Pope and Medzhitov, 2018). Our gSWITCH web application is a powerful tool for identifying specific dynamic patterns based on progressive changes in gene expression during cellular state transitions or any variable-dependent changes. This can be particularly valuable for discovering novel mechanisms in defined biological contexts. Switch genes signify the neurogenic commitment of cortical progenitors and have enabled the identification of new regulators of brain corticogenesis (Aprea et al., 2013). However, no biologist-friendly tool has been available to systematically detect such genes. gSWITCH is a biologically inspired method, and here we experimentally validate its prediction.

Using gSWITCH and following a data driven approach, we identified Bcl11a as a critical switch gene in the transition from OPCs to pre-OLs. To test this prediction, we performed loss-of-function (LOF) experiments using siRNA and found that inhibition of Bcl11a impaired OPC differentiation into oligodendrocytes. Because Bcl11a expression is markedly down-regulated in aged OPCs, we next carried out gain-of-function (GOF) experiments both *in vitro* and *in vivo*. Restoring Bcl11a expression in aged OPCs significantly enhanced their differentiation. Together, the LOF and GOF experiments unambiguously validate the key role of Bcl11a in oligodendrocyte lineage progression and support the gSWITCH prediction.

The extent to which the reduced differentiation capacity of aged OPCs is intrinsically encoded within the cells themselves or induced by prolonged exposure to an aged tissue environment is an interesting question. Based on our previous work, we favour the view that loss of OPC function is primarily determined extrinsically since various manipulations of the aged environment such as heterochronic parabiosis (Ruckh et al., 2012), fasting and calorie restriction mimetics (Neumann et al., 2019), and niche biomechanics (Segel et al., 2019) can all alter the cell-intrinsic state, reverting aged cells to a ‘youthful state’. Significantly, when aged OPCs are transplanted into the neonatal CNS they proliferate and differentiate as if they were neonatal OPCs (Segel et al., 2019). The reversion of aged OPCs to a functional state by changes in their external environment necessarily operates through changes in cell-intrinsic function, suggesting that the same intrinsic mechanisms could be targeted directly to restore declining OPC function—for example through epigenetic regulation of differentiation inhibitors (Shen et al., 2008) or overexpression of transcriptional regulators such as c-Myc (Neumann et al., 2021, Dimas et al., 2025).

Ageing is undeniably complex, with multiple cellular, metabolic, and extracellular changes, inflammation occurring in parallel. Discovering new regulators remains essential for understanding how these layers collectively shape age-related decline. Interventions used so far have shown that no single mechanism can fully explain the functional deterioration of ageing oligodendrocyte progenitors. Instead, each factor highlights a distinct point at which the ageing process exerts pressure, gradually revealing the distributed architecture through which ageing unfolds. Within this landscape, our identification of Bcl11a—emerging from an analysis of oligodendrocyte lineage progression—defines another biologically meaningful regulatory axis involved in these age-associated changes, expanding the mechanistic resolution with which ageing can be understood. Only by charting these individual components can we explain how diverse age-related diseases emerge and begin to design personalised strategies capable of addressing the true complexity of ageing in a mechanistically grounded way.

BCL11A is a zinc-finger transcription factor primarily known as a transcriptional repressor, though it can also function as an activator (Horton et al., 2025; Bauer and Orkin 2015; Lee et al., 2017). Our computational molecular-evolution analyses indicate that Bcl11a is a vertebrate-specific transcription factor that has undergone pervasive purifying selection. In this context, Bcl11a is perhaps evolutionarily linked to major cellular programs, and its dysfunction may contribute to disease. In haemoglobin regulation, Bcl11a orchestrates the fetal-to-adult haemoglobin switch by binding to and repressing the γ-globin genes, thereby enabling activation of adult β-globin—a critical transition that supports the oxygen-transport requirements of postnatal life (Liu et al., 2018). In oligodendrocytes, we find that Bcl11a is essential for lineage progression. Bcl11a expression in young OPCs is important for maintaining their functional resilience, ensuring they retain a strong capacity to differentiate efficiently.

Notably, Bcl11a deficiency or dysfunction has been implicated in autism spectrum disorder and intellectual disability based on human genetic studies (De Rubeis et al., 2014; Dias t al., 2016; Cai et al., 2017). In the hematopoietic system, Bcl11a deficiency in hematopoietic stem cells induces an ageing-like state (Luc et al., 2016). Consistent with this, we observed that aged OPCs, despite being impaired in their differentiation capacity, were able to regain functional resilience when Bcl11a expression was restored. In this sense, Bcl11a reinstates functional robustness in aged OPCs. Future studies will elucidate the detailed gene-regulatory mechanisms through which Bcl11a functions in young OPCs, clarify how it enhances functional competence in aged OPCs, and further define its potential as a therapeutic target.

## Materials and Method

### gSWITCH

[This includes section g.1 to g.5]

**g.1 Usage:**

It is designed to handle gene-expression data from lineage progression or time series or any variable dependent experiments. Experiment type: bulk RNA-seq, single cell RNA-seq experiments flattened to pseudo bulk. In the series data, the number of series or time points or group ≥3 and the replicates per time point or group≥2.

g.2 **Access**: freely available at https://altoslabs.shinyapps.io/gSWITCH/

**g.3 Implementation**

gSWITCH is a web application and operating system (OS) independent. gSWITCH is implemented in ‘R’ by using *reshape2*, *dplyr*, *DEseq2*, *ggplot2*, *tximport*, *MonoInc* and *plotly* (R Core Team, 2019; Wickham, 2007; Wickham, 2016; Wickham, 2017, Love et al., 2014) and employing RStudio’s *shiny* and *shinycssloaders* packages for web application.

**g.4 Logic**

gSWITCH is based on the following: 1) GLM (Generalised linear model) fit and hypothesis testing, 2) specific contrasts analysis, 3) absolute maximum/minimum (in a closed interval) and monotonicity to categorize switch types.

We refer the **Supplementary Figure 2**. We employ DESeq2 for GLM fit, calculation of LTR (likelihood ratio test) p-value (adjusted by FDR) and select genes for which adjusted p value (Adj p) < user-defined-value (u.d.) [0.05 is set as default]. Let say these genes are our set (i). We then calculate contrast between the first and the last cell state or time points and the genes for which Adj p < user defined (or 0.05 as default) and the magnitude of fold change |FC| > user-defined-value (or 1.4 is set as default) was considered as set (ii). Intersection of (i) and (ii) was taken to identify four different switch types which are based on absolute max/min and monotonicity. A relative expression value (g) was calculated by determining expression level of each cell state relative to the mid-point [for n=odd number: mid-point={(n/2)+0.5]}^th^ cell state; for n=even number: mid-point=(n/2)^th^ cell state].

1. Type 1: properties: (a) absolute maxima at the end point (b) ‘g’ is monotonically increasing i.e. within [g1, g_n_], g_j-1_<g_j_ (j=1,2,…n) n is a positive integer ≥ 3 (c) contrast analysis: X_n_ vs X_n-1_: Adj p < u.d. (or 0.05), |FC| > 1.4 (or u.d.).
2. Type 2: properties: (a) absolute maxima at the mid-point (b) ‘g’ is monotonically increasing from start to mid-point and then monotonically decreasing from the mid-point to end point i.e. within [g_1_,g_j_], g_j_ ≥ g_j-1_ AND [g_j_,g_n_], g_j_ ≥ g_j+1_ (c) contrast analyses: X_j_ vs X_j+1_ AND X_j-1_ vs X_j_ : Adj p<0.05 (or u. d.)], |FC| > 1.4 (or u.d.).
3. Type 3: properties: (a) absolute maxima at the start point (b) ‘g’ is monotonically decreasing as it progresses to end point i.e. within [g_1_, g_n_], g_j-1_ ≥ g_j_ (c) contrast analysis: X_1_ vs X_2_ : Adj p<0.05 (or u.d.), |FC| >1.4 (or u.d.).
4. Type 4: properties: (a) absolute minima at the mid-point (b) ‘g’ is monotonically decreasing from start to the mid-point and then monotinically increasing from the mid-point to the end point i.e. within [g_1_,g_j_], g_j-1_ ≥ g_j_ AND [g_j_,g_n_], g_j_ ≤ g_j+1_. (c) contrast analyses: X_j_ vs X_j+1_ AND X_j-1_ vs X_j_ : Adj p<0.05 (or user defined)], |FC| >1.4 (or u.d.).

Identifying transcription factors (TFs)

Comprehensive list of TFs were retrieved from TFcheckpoint database (Chawla K et al., 2013), an experimental evidence-based database. L∩S is the TFs in the switch genes. L: list from TFcheckpoint, S: switch genes.

**g.5 Parameters used in this paper**

Adj p<0.05, FC=1.5

### RNAseq analysis

RNA was isolated using RNA MiniPrep (Zymo Research, D7001) following manufacturer protocol. Dnase treatment on column (as per manufacturer protocol) was included in this isolation protocol. Paired end directional library was prepared using NuGen kit: Ovation RNA-seq Systems 1-16 for model organisms (0349-32).

Oligodendrocyte lineage dataset: The accession number of this dataset is **GSE301578**. Data were pre-processed through sortMeRNA, trimmomatics. Rat transcriptome (Rnor_6.0.cdna+Rnor_6.0.ncrna) index was build and quantification was performed using Salmon (Quasi mode). Data was normalised using DEseq2. This tximport of salmon quantification were run on gSWITCH (accessible at https://altoslabs.shinyapps.io/gSWITCH/) for switch genes identification and visualisation.

Analysis of aged and young Rat OPC data: The accession number of this dataset is **GSE303317**. Data were pre-processed using sortMeRNA and Trimmomatic. The rat transcriptome index (Rnor_6.0.cdna + Rnor_6.0.ncrna) was built, and quantification was performed using Salmon in quasi-mapping mode. Raw Salmon counts were imported using the tximport package. Differential expression analysis was conducted with DESeq2, incorporating batch as a covariate in the design formula: design = ∼ batch + group. P-values were adjusted for multiple testing using the false discovery rate (FDR) method. Genes with an adjusted p-value less than 0.05 and a fold change greater than or equal to 1.5 were considered differentially expressed between the two groups. For data visualization, counts were normalized using a variance stabilizing transformation (VST) (varianceStabilizingTransformation function from DESeq2), and batch effects were removed using the removeBatchEffect function from the limma package (Ritchie et al., 2015).

### TF-binding site enrichment analysis

Using type 3 and type 4 switches as input, we employed the ChIP-X Enrichment Analysis (ChEA3) tool [https://maayanlab.cloud/chea3/] (Keenan et al., 2019) and utilised two experimentally curated ChIP-seq databases—ENCODE (Encyclopedia of DNA Elements) (ENCODE Project Consortium, 2012) and ReMap2 (Regulatory Map database) (Hammal et al., 2022) —to identify transcription factors (TFs) with significantly enriched binding sites (FDR < 0.05). We then focused on transcription factors consistently identified by both ENCODE and ReMap2.

### dN/dS analysis

Twenty-three vertebrate species (including lamprey), spanning a broad range of vertebrate phyla, were used in this study (**Supplementary Table 4**). The longest (XL) coding sequences (CDS) and the corresponding protein sequences of Bcl11a were obtained from Ensembl (Dyer et al., 2025). For lamprey, the sequences were retrieved from GenBank (Sayers et al., 2021), as they were not available in Ensembl. Protein sequences were aligned using MAFFT (Katoh and Standley, 2013), and the resulting alignment together with the corresponding coding sequences was used as input to the PAL2NAL program (Suyama et al., 2006) to generate codon-based nucleotide alignments. The species tree was generated from CDS using the NGPhylogeny server (Lemoine et al., 2019), the nucleotide sequence alignment, BMGE (Criscuolo and Gribaldo, 2010) for alignment curation, and PhyML (Guindon et al., 2010) for tree inference. dN/dS ratios and likelihood ratio tests of model comparisons were performed using Codeml in the PAML package v4.8a (Yang, 2007). Codon frequency was set to F3X4, the universal genetic code was specified in the icode variable, and the program was run with the following selection model parameters: Nssites 0 and 3 (selection models M0 and M3), Nssites 1 and 2 (models M1a and M2a), and Nssites 7 and 8 (models M7 and M8). Significance of log-likelihood differences for model comparisons was tested using χ^2^ statistics with appropriate degrees of freedom, as described in the PAML manual (https://github.com/abacus-gene/paml/blob/master/doc/pamlDOC.pdf).

Further detail on the M7 and M8 comparison: these models differ only by M8 allowing an additional class of sites with dN/dS > 1. The likelihood ratio test (LRT) comparing these indicates a significant preference for M8 (P = 0.006) but identifies only a single site with posterior mean dN/dS > 1 and confirms all other sites to be evolving under purifying selection. (Codon 156, i.e. nucleotide positions 466–468 in our curated alignment, is inferred to have dN/dS = 2.93 ± 2.81 and posterior probability 0.94 of being under positive selection. However, inspection of the alignment reveals that this position is represented only in macaque, rabbit, blue tit, and frog, making evidence for positive selection across species unconvincing even at this one site.)

### Animals

Sprague–Dawley rats (C. River, Margate, UK) and C57Bl/6 mice (18-month-old) were housed in individually ventilated cages at 22 °C ± 1 °C and 60% ± 5% humidity under a 12-h light–dark cycle. Food and water were provided ad libitum in a standard rodent facility at the University of Cambridge, UK or Altos Labs-Cambridge Institute of Science, UK. All procedures were carried out in accordance with the Animals (Scientific Procedures) Act 1986 (Amendment Regulations 2012) and were approved by the Animal Welfare and Ethical Review Body (AWERB) of the University of Cambridge and Agenda Resource Management.

### Cellular isolation

Rats were sacrificed using an overdose of pentobarbital and the whole brain was dissected immediately in ice cold hibernate A low fluorescence (HALF) medium (PMID: 31585093). Subsequently cellular isolation was performed as described. Per 10 million cell suspension, we used 2.5 μg of primary antibody. Oligodendrocytes (OL) were isolated first using goat-anti-mouse-MOG-Biotinylated (RD systems, BAF2439); followed by OPC were isolated from the MOG negative population using mouse-anti-rat-A2B5-IgM antibody (Millipore; MAB312). Finally, pre oligodendrocytes (PreOL) were isolated from MOG and A2B5 negative population using mouse-anti-O4-IgM (R&D systems, MAB1326). The respective secondary antibodies (20 ul) were: mouse-anti-biotin magnetic micro beads (Miltenyi, 130-105-637), rat-anti-mouse-IgM antibody magnetic beads (Miltenyi, 130-047-302), rat-anti-mouse-IgM magnetic beads (Miltenyi; cat#130-047-302).

### Cell culture and differentiation

Culture of aged OPCs (isolated from 20-24-month-old rats) were performed as described (PMID: 31585093). Additionally, DMEM-F12 (Sigma), GlutaMAX^TM^ (Gibco^TM^) and B27 supplement (Gibco^TM^) has been used in the media for aged OPC culture. Proliferation medium contains PDGF and bFGF at 30 ng/ml each (PeproTech). For differentiation as described (Ghosh et al., 2024) OPC medium was supplemented with 40 ng ml^-1^ 3,30,5-Triiodo-L-thyronine (T3) (Sigma; T6397). Medium was replaced on alternate days for 5 days.

Neonatal OPCs (isolated from P7 rats) were cultured as described (Ghosh T et al., 2024).

### Cloning of Bcl11a

The original Human XL BCL11A clone was a gift from Prof. Walid Khaled lab (Cambridge University, UK). BCL11A was subcloned to a TOPO vector (pCR™ 2.1-TOPO™ TA using TOPO™ TA Cloning™ Kit following manufacturer instruction) and sequence was verified. For *In vitro* transcription, the following primers were used for PCR amplification of BCL11A:

FP: 5’ *TAATACGACTCACTATAG*gccaccATGTCTCGCCGCAAGCAA 3’

(italics: T7 promoter sequence, small case: kozac sequence)

RP: 5’ CTATTCAGTTTTTATATCATTATTCAACACTCGATCA 3’

Purified PCR product (from above) was used as a template and transcribed *in vitro* using HiScribe^TM^ T7 ARCA mRNA Kit (New England Biolabs) with tailing and modified nucleotides (Tebu Bioscience) following manufacturer protocol (E2060S). mRNA was purified using Directzol^TM^ RNA Miniprep kit (Zymo Research). CleanCap^R^ EGFP mRNA (L-7601) was procured from Tebu Bioscience.

### Western blot analysis

MCF7 cells were used to verify protein-level expression of BCL11A following transfection with in-vitro–transcribed Bcl11a mRNA (see above). MCF7 cells were selected because they exhibit poor endogenous BCL11A expression (Waterhouse M et al., 2025). For mRNA transfection method (see below). After 36 hours of transfection, cells were lysed in the wells using RIPA buffer containing protease inhibitors. The lysates were incubated at 4 °C for 30 min and centrifuged at maximum speed at 4 °C for 30 min. The resulting supernatant was used for protein separation on a 7.5% SDS–PAGE gel and subsequently transferred to a PVDF membrane for Western blotting. Antibodies used were, anti-BCL11A (ab191401, Abcam, 1:3000) (Waterhouse et al., 2025) and an HRP-conjugated donkey anti-rabbit IgG secondary (NA934, Amersham/GE Healthcare) (1:10000); anti-beta Actin antibody (ab6276) (1:3000) and HRP-conjugated sheep anti-mouse IgG secondary (NA931, Amersham/GE Healthcare) (1:10000). Immunoreactive bands were detected using ECL reagent (Amersham) and imaged an Odyssey Imaging system.

### Transfection

siRNA transfection in neonatal OPCs: We used Lipofectamine^TM^ RNAiMAX transfection reagent (ThermoFisher scientific) and follow the protocol as described (Ghosh et al., 2024). siRNAs were procured from Dharmacon^TM^. For Bcl11a we used siGENOME Rat Bcl11a (305589)-SMARTpool (Cat no. M-085668-01-0010); for control we used siGENOME non-Targeting siRNA Pool #2 (Cat no. D-001206-14-05)

mRNA transfection in aged OPCs: 60,000 cells per well of a 24 well plate was seeded. Cells were kept in proliferation media [PDGF and bFGF, 30ng/ml each (Peprotech)] for 5 days. mRNA (approx. 500ng) was transfected using Lipofectamine^TM^ MessengerMax^TM^ transfection reagent (ThermoFisher scientific) following manufacturer protocol.

### Immunofluorescence

Cells were fixed in 4% paraformaldehyde (PFA) for 10 min at room temperature, washed in PBS, and processed for immunofluorescence staining as described previously (Neumann et al., 2019). For in vivo experiments, animals were perfused with 4% PFA in PBS; spinal cords were post-fixed overnight in 4% PFA, cryoprotected in 20% sucrose, and embedded in OCT medium (Tissue-Tek). Immunofluorescence staining was performed on 12 µm cryostat sections following previously described method (Rawji et al., 2018).

The antibodies used in this study, described previously (PMID: 38364788), were: rat anti-MBP (MCA409S, Serotec) [1:500], rabbit anti-OLIG2 (AB9610, Millipore) [1:1000 or 1:100], chicken anti-GFP (ab13970, Abcam) [1:1000], donkey CyTM3 AffiniPure anti-Rat (712-165-150, Jackson Immuno Research) [1:1000], donkey Alexa Fluor 647 anti-Rabbit (A31573, Invitrogen) [1:1000 or 1:500], donkey Alexa Fluor 488 Anti-Chicken (703-546-155, Jackson Immuno Research) [1:1000].

Images were acquired using SP8 (Leica TCS) confocal microscope. All acquisition settings were kept constant throughout the experiment. Images were processed in ImageJ version 2.1.0 (Schindelin et al., 2012). Cell counting was performed using the Cell Counter tool in ImageJ. Immunofluorescence experiments were performed and analysed blinded to treatment.

### EdU proliferation assay

For EdU incorporation, cells were incubated with 10 µM EdU for 5 hours and subsequently processed following the manufacturer’s instructions (Click-iT Plus EdU Alexa Fluor 647 Imaging Kit, Thermo Fisher, C10640). In vitro tissue-culture cells were fixed in 4% paraformaldehyde (Thermo Fisher, 10131580) for 10 minutes at room temperature. Cells were washed once with PBS and blocked in PBS containing 0.1% Triton X-100 and 5% donkey serum (Sigma, D9663) for 30 minutes at room temperature. EdU detection was performed according to the kit protocol, followed by immunostaining. Antibodies used were: Anti-OLIG2: rabbit anti-OLIG2 (AB9610, Millipore) [1:1000], Goat Alexa Fluor 488 anti-rabbit IgG (A11008, Invitrogen) (1:1000) [siRNA experiments in Figure 4), Goat Alexa Fluor 594 anti-rabbit IgG (A11012, Invitrogen) (1:1000) [mRNA transfection experiment in Figure 5].

### RT-qPCR

Total RNA was isolated using the mirVana kit (Ambion). RNA samples were treated with DNase I (Thermo Fisher Scientific) prior to reverse transcription. cDNA was generated using random hexamers (Invitrogen) and SuperScript IV Reverse Transcriptase (Invitrogen) according to the manufacturer’s protocol. Real-time PCR was performed using SYBR Green Master Mix (Applied Biosystems) on a QuantStudio 7 Flex system (Applied Biosystems). The PCR program was: 50 °C for 2 min, 95 °C for 10 min, followed by 40 cycles of 95 °C for 15 s and 60 °C for 1 min. A melt-curve (dissociation) analysis was performed at the end.

Relative quantification of qPCR data was performed using the Pfaffl method (Pfaffl MW, 2001). PCR efficiencies for the target gene (Bcl11a) and the endogenous control (Actb) were determined from the slopes of their respective standard curves. For qPCR performed presented in Supplementary Figure 4, Hprt was used as endogenous control for normalisation.

Rat specific primers:

Bcl11a:

F: GTGACACCAGAAGATGACGA

R: ATTAGGGCTCCGTGTGCAG

[Amplicon: 131 bp] Actb:

F: CACCATGTACCCAGGCATTG

R: CACACAGAGTACTTGCGCTC

[Amplicon: 111 bp] Hprt:

F: CAGACTTTGCTTTCCTTGGTCA

R: CCAACACTTCGAGAGGTCCT

[Amplicon: 89 bp]

Primer sequences for Pdgfra and Mbp were as described previously in Ghosh et al. (2024).

### AAV delivery

The SOX10 multiple-species conserved enhancer element 5, conjugated to the *cfos* basal promoter (Addgene #115783; Pol et al., 2013), was used to drive oligodendrocyte-lineage–specific expression of Bcl11a and EGfp (**Figure 6A**). For the control construct, the same regulatory sequence drove expression of EGfp alone. The entire SOX10-driven expression cassette (**Figure 6A**) was cloned between the inverted terminal repeats (ITRs) of an adeno-associated virus 2 (AAV2) vector. Cloning, large-scale AAV-PHP.eB packaging, and purification were performed by GenScript (UK).

AAV carrying the construct (**Figure 6A**) was administered to 18-month-old mice via tail vein injection at a dose of 5 × 10¹¹ genome copies (GC) in a total volume of 200 µL.

### Focal demyelination of the spinal cord

A mouse spinal cord demyelination model was generated by injecting 1% lysophosphatidylcholine (lysolecithin, LPC; Sigma-Aldrich, Gillingham, UK) prepared in PBS into the ventral white matter of the spinal cord at the Th13/L1 level following surgical exposure under isoflurane anaesthesia. 18-month-old C57Bl/6 mice were used in this study; the animals do not manifest abnormalities in this model. Mice were sacrificed by perfusion fixation at 14 days post lesioning (dpl) for histological analysis.

### Plp1 *in situ* hybridisations (ISH)

We followed standard procedures used in the Richardson laboratory (UCL) (https://www.ucl.ac.uk/~ucbzwdr/In%20Situ%20Protocol.pdf). Plp1 mRNA was detected using a digoxigenin (DIG)-labelled riboprobe. Primers (FP: GGGGATGCCTGAGAAGGT; RP: TGTGATGCTTTCTGCCCA) targeting the Allen Brain Atlas Plp1 sequence (ABA experiment 75496148) were used to amplify the region, which was subsequently TA-cloned into the pCRII vector using the Thermo Fisher K206001 kit. Insert orientation was confirmed by Sanger sequencing. Following linearisation with XbaI, an antisense DIG-labelled riboprobe (970 bp) was synthesised by in vitro transcription using T7 RNA polymerase.

For hybridisation, the DIG-labelled probe was diluted 1:1000 in hybridisation buffer [50% de-ionised formamide, 1× salts (0.2 M NaCl, 5 mM EDTA, 10 mM Tris-HCl pH 7.5, 5 mM NaH₂PO₄, 5 mM Na₂HPO₄), 0.1 mg/ml yeast tRNA, 10% dextran sulphate, and 1× Denhardt’s solution], denatured at 70–75 °C for 10 min, and applied to tissue sections. Slides were incubated overnight at 65 °C in a humidified chamber containing 50% formamide/1X SSC. The next day, sections were washed in three pre-warmed solutions of 50% formamide/1X SSC/0.1% Tween-20 at 65 °C, followed by two room-temperature washes in 1X MABT (100 mM maleic acid, 150 mM NaCl, 0.1% Tween-20, pH 7.5). Sections were blocked for 1 h in 20% sheep serum and 5X blocking reagent (2% Roche blocking reagent + 10% heat-inactivated sheep serum) in 1X MABT, then incubated overnight at 4 °C with anti-DIG-AP antibody (1:1000) prepared in the same blocking solution. After three washes in MABT, colour development was performed using NBT/BCIP in a freshly prepared Tris/MgCl₂/NaCl developing buffer with 5% polyvinyl alcohol (PVA) at pH 9.8, incubated at 37 °C in the dark until optimal signal intensity was achieved. Slides were rinsed in PBS, counterstained with DAPI, and mounted in Fluoromount™ (Sigma) and imaged under Thermo Fisher EVOS M5000 Cell Imaging System. ISH experiments were performed and analysed blinded to treatment.

### Statistical analysis

R (version 4.0.2) or GraphPad Prism (version 8.4.3) was used for statistical analyses. Normality was evaluated using the Shapiro–Wilk test, and Welch’s t-test was used in cases of heteroscedasticity.

## Supporting information

Switches in OL lineage

Transcription factor (TF) switches in OL lineage

## Acknowledgements

This work was supported by grants from Adelson Medical Research Foundation (AMRF) (R.J.M.F.), UK Multiple Sclerosis Society grant MS50 (R.J.M.F.), Wellcome Trust-Medical Research Council Cambridge Stem Cell Institute core support grant 203151/Z/16/Z (R.J.M.F.), Altos Labs- Cambridge Institute of Science (R.J.M.F. and T.G.), European Molecular Biology Laboratory (EMBL) (N.G.). We thank Dr. Maike Paramor at the NGS Library Facility at the Cambridge Stem Cell Institute for RNA sequencing and advice, Prof. Walid Khaled and Dr. Mark Waterhouse for providing the Bcl11a clone, and Prof. Daniel Geschwind and Dr. Riki Kawaguchi for helpful discussions. We thank Prof. Gonçalo Castelo-Branco and Dr. Eneritz Agirre for kindly sharing Bcl11a expression fold-change comparing myelinating oligodendrocytes with OPCs from their published single-cell RNA-seq dataset.

## Author contributions

Conceptualization: T.G. and R.J.M.F.; formal analysis: T.G.; developer of gSWITCH application: T.G., Computational molecular evolution analysis: T.G. and N.G.; investigation: T.G., R.B., C.Z., A.S., W.H.A., A.L.; methodology: T.G., R.B., C.Z., AS, N.G., R.J.M.F.; project administration: T.G. and R.J.M.F.; supervision: R.J.M.F. and T.G.; writing – original draft, T.G. and R.J.M.F.; writing – review & editing, T.G., R.J.M.F., N.G., R.B., C.Z., A.S., W.H.A., A.L.

## Data availability

GEO accession number of RNA-seq datasets: GSE301578, GSE303317.

## Figures and corresponding Legends

**Supplementary Figure 1.**
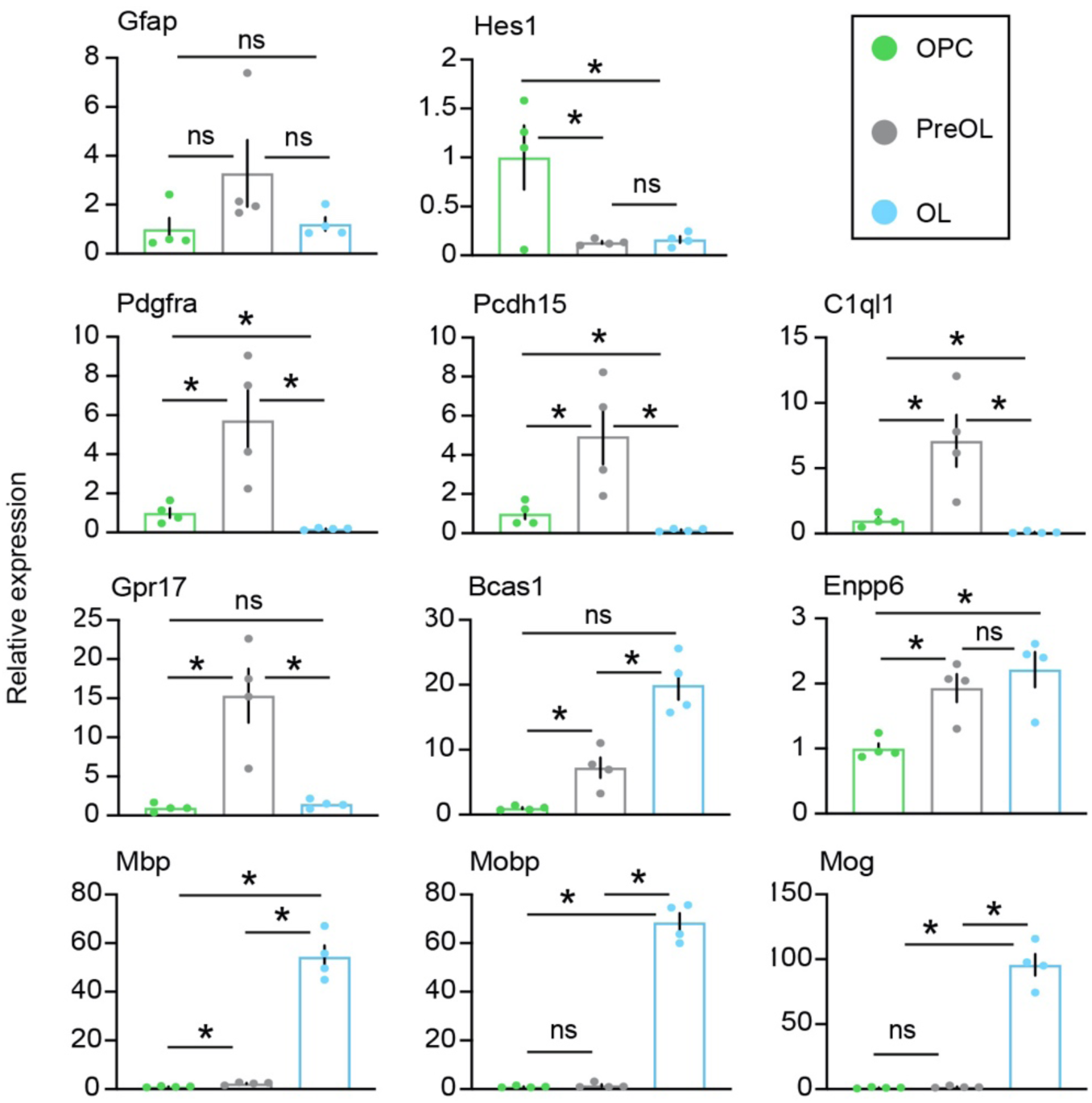
Normalised expression values of marker genes (for astrocytes, progenitor-state marker, OPC, PreOL, and OL) from RNA-seq data (GEO accession number: GSE301578) were plotted relative to OPCs. n=4 (young rats), mean±SEM, * adjusted P<0.05, Benjamini–Hochberg FDR after Wald test. See also Supplementary text.

**Supplementary Figure 2.**
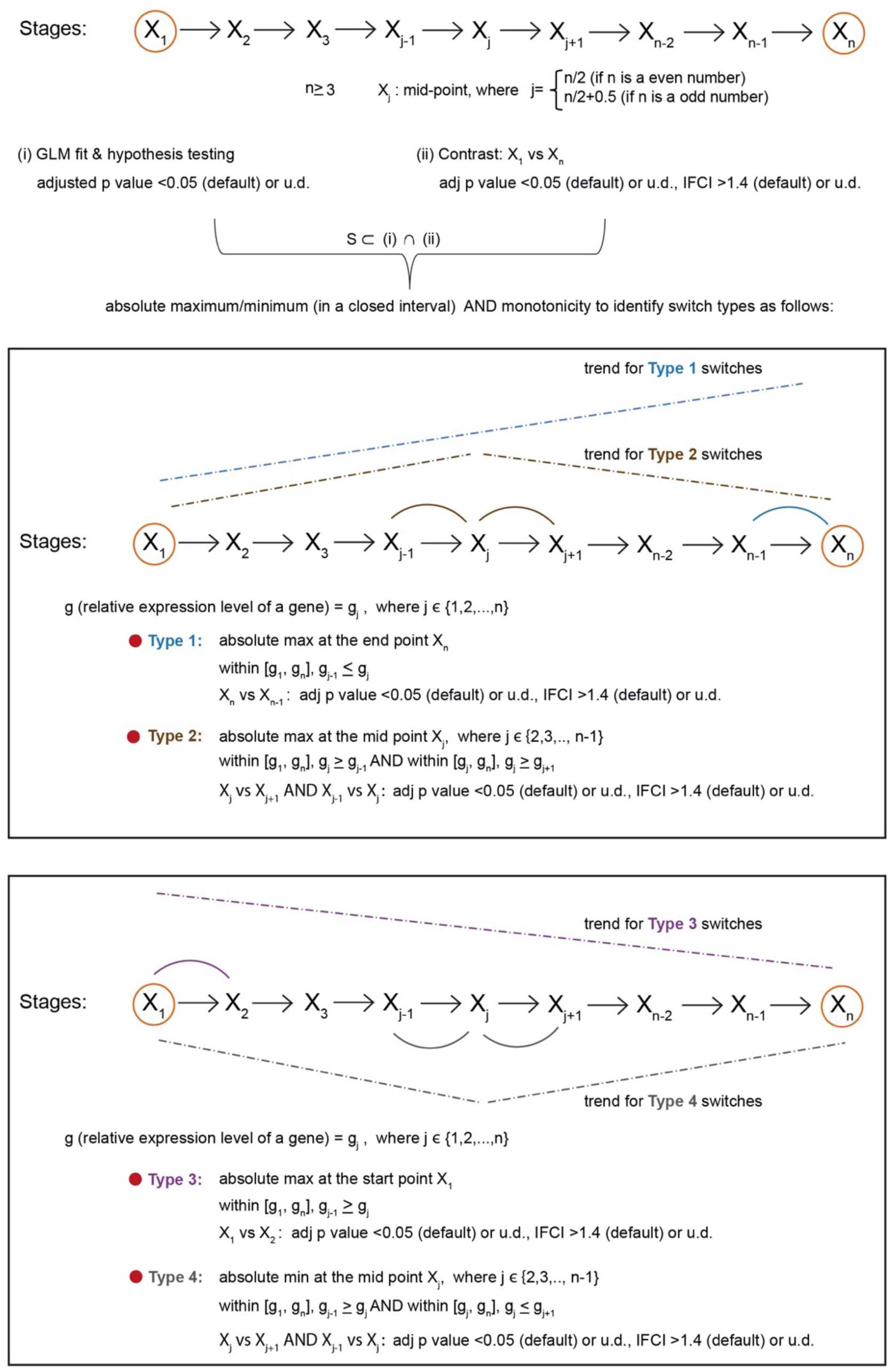
Schematic representation of the logic of the gSWITCH web application. (for detail see ‘*Materials and Methods*’ section). The gSWITCH application is accessible at https://altoslabs.shinyapps.io/gSWITCH/. Red circles indicate the start and the end state of a series (time or lineage stages or any other variable dependent series). GLM fit and hypothesis testing followed by contrast analysis between start and the end state, generates a subset of genes on which we apply maximum (max) or minimum (min) criteria within a closed interval while taking account of monotonicity trend as well as further specific contrasts between groups. This allows identification of four types of switches. Detail criteria are shown inside the boxes and the trend pattern are represented by coloured dashed-straight lines. Arc or solid curve lines indicate statistical significance for specific contrast. Adj p value: Adjusted p value [Benjamini–Hochberg FDR after LRT or Benjamini–Hochberg FDR after Wald test for specific contrast analysis]. u.d.: user defined value. Note that to run gSWITCH, RNA-seq data are required (bulk or scRNA-seq flattened to pseudo bulk), where the number of series or time points or group ≥3 and the replicates per time point or group≥2. gSWITCH also identifies transcription factors (TFs) within the switches and provides a summary of the total number of switches and TFs per switch type.

**Supplementary Figure 3.**
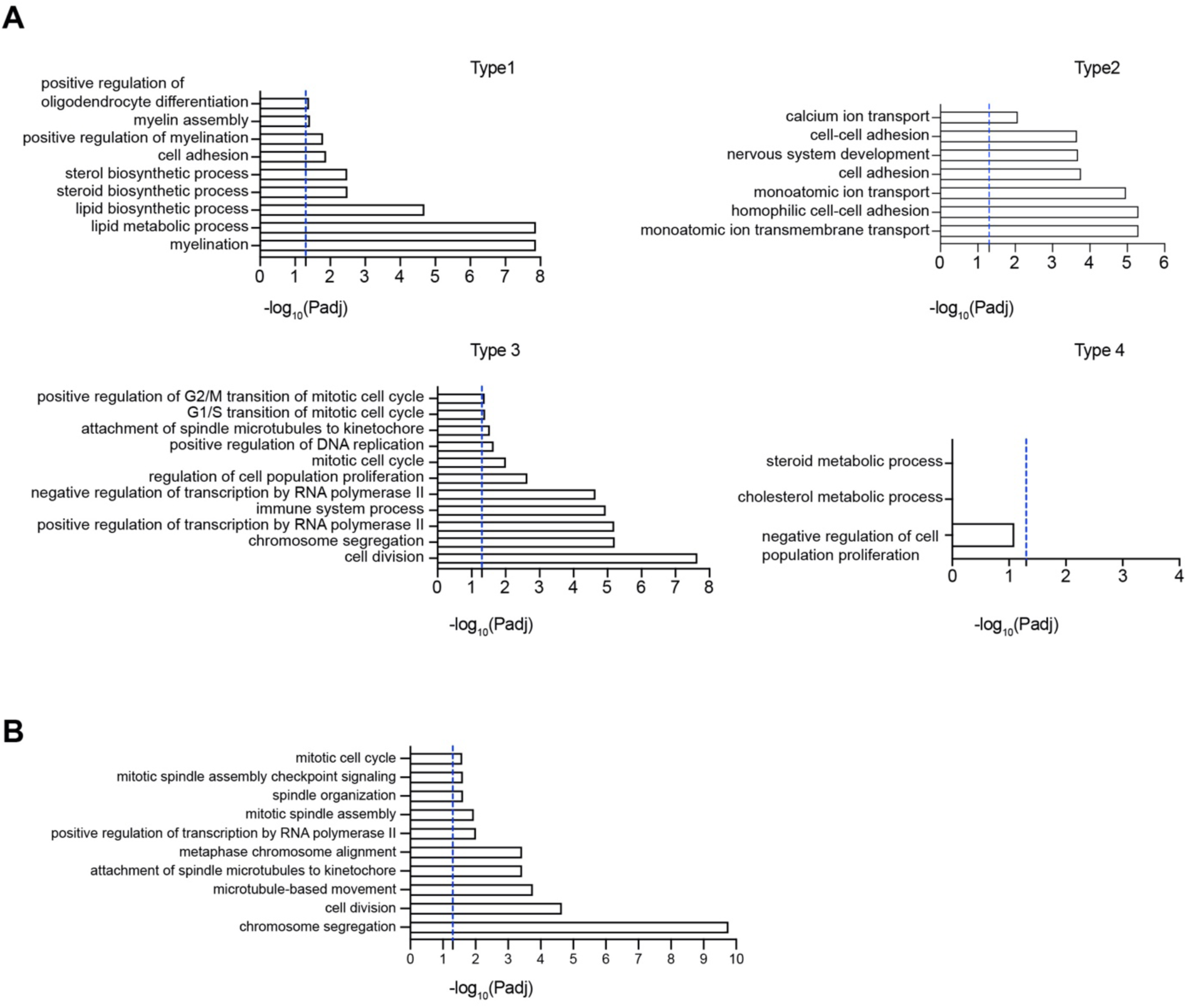
Gene ontology (GO) analysis. **A)** Top over-represented GO terms for each switch type shown in Figure 1C. **B)** Top over-represented GO terms for the 135 overlapping genes shown in Figure 2C, defined as genes that belong to Type 3 switches and are also affected by ageing. Padj: Benjamini–Hochberg adjusted P value. The dotted blue line indicates the significance threshold of Padj = 0.05, corresponding to −log10(Padj) = 1.3. Please note that only 8 Type 4 genes overlapped with differentially expressed genes in ageing. Due to this small number, we could not identify any significant GO-term enrichment, and therefore this was not plotted.

**Supplementary Figure 4.**
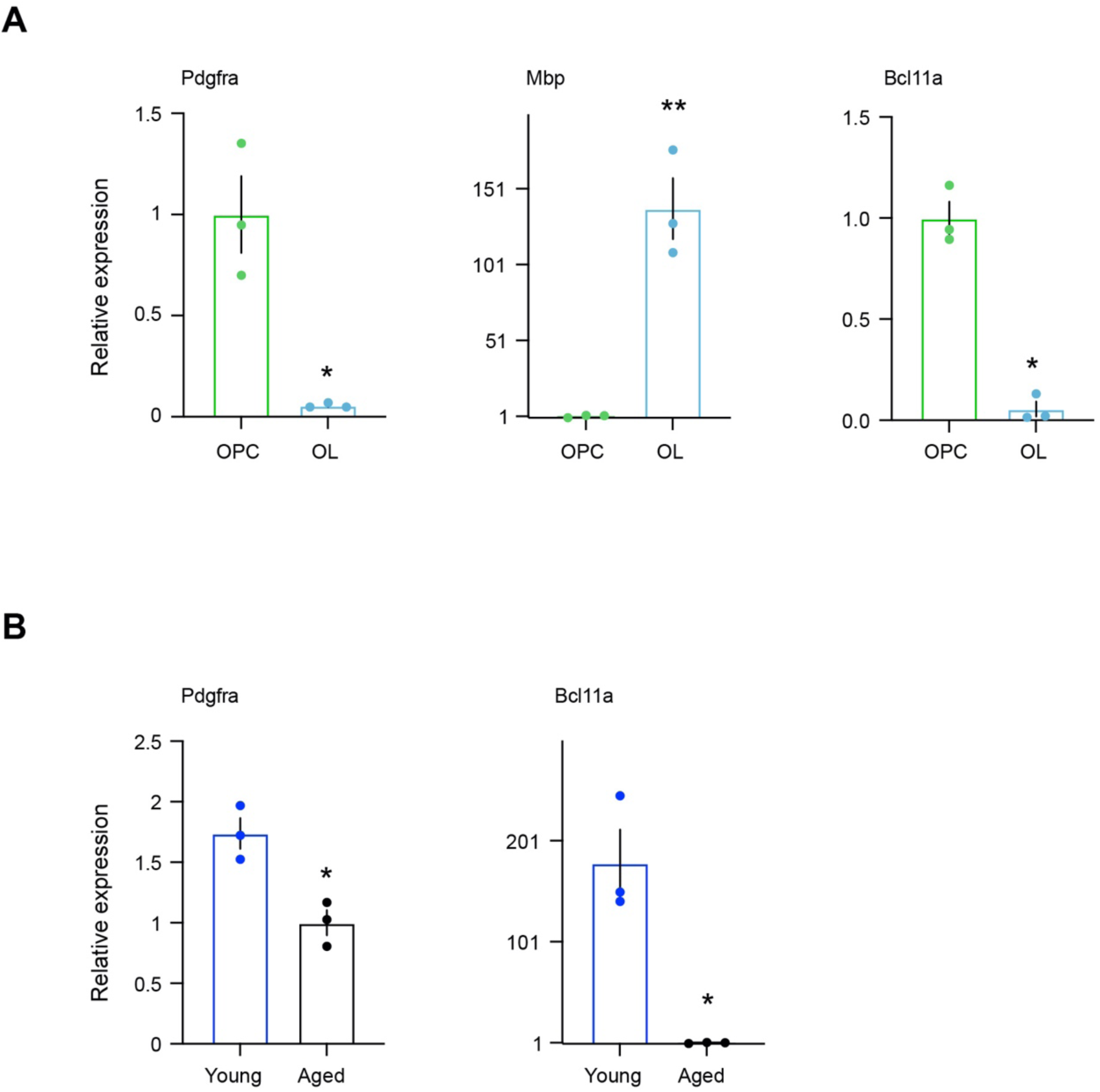
Bcl11a expression in OPCs, OLs and aged OPCs, as determined by RT-qPCR (reverse transcription followed by qPCR). **A)** Expression levels of Bcl11a, Pdgfra as an OPC marker, and Mbp as an OL marker were measured in OPCs and OLs and plotted relative to OPCs. **B)** Expression levels of Bcl11a and Pdgfra were measured in young and aged OPCs and plotted relative to aged OPCs. **(A-B)** n=3 rats, mean±SEM, * Bonferroni adjusted P<0.05, ** Bonferroni adjusted P<0.01, Student’s t test (unpaired, two tailed).

**Supplementary Figure 5.**
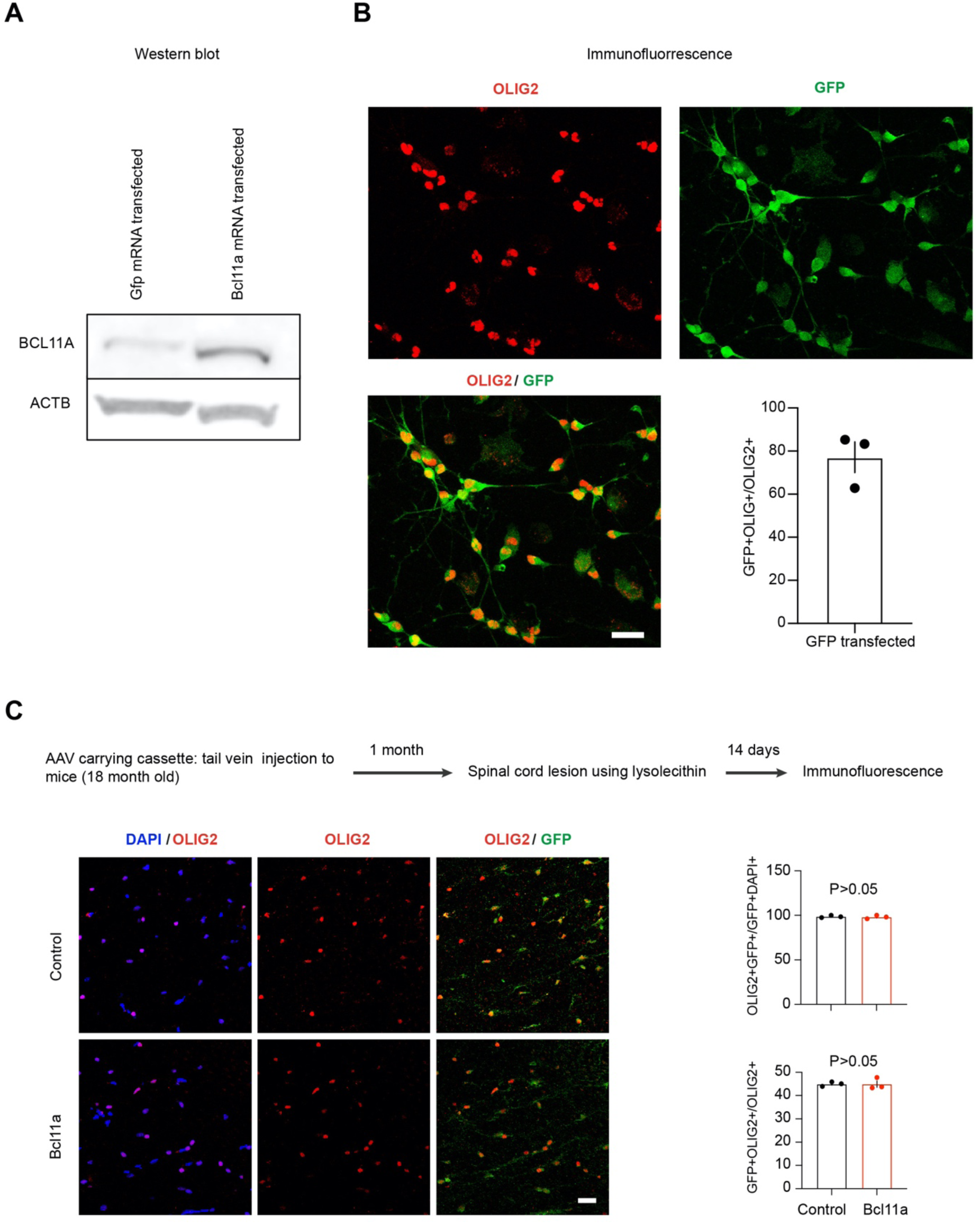
Fidelity of protein expression from in vitro–transcribed Bcl11a mRNA; mRNA transfection efficiency in aged OPCs; and fidelity of AAV construct expression in the OL lineage and infection efficiency in aged mouse brain. **A)** Western blot analysis using antibodies to BCL11A and β-actin (ACTB), with ACTB serving as a loading control, in MCF7 cells transfected with Bcl11a mRNA or Gfp mRNA (control). **B)** Immunofluorescence analysis using antibodies to OLIG2 and GFP in aged OPCs transfected with Gfp mRNA. Quantification of GFP+OLIG2+ cells were plotted as percentage of OLIG2+ cells. n = 3 independent experiments (each using OPCs isolated from pooled brains of 2 aged rats). mean ± SEM. Scale bar: 22 µm. **C)** AAV carrying constructs are shown in **Figure 6A**. Left: Immunofluorescence analysis of OLIG2 and GFP immunostaining. Right top: Quantification of OLIG2+GFP+ cells were plotted as a percentage of GFP+DAPI+ cells. Right bottom: quantification of OLIG2+GFP+ cells were plotted as a percentage to OLIG2+ cells. n= 3 mice, mean+SEM. P: p-value, Student’s t test (unpaired, two tailed). Scale bar: 20 µm.

**Supplementary Figure 6.**
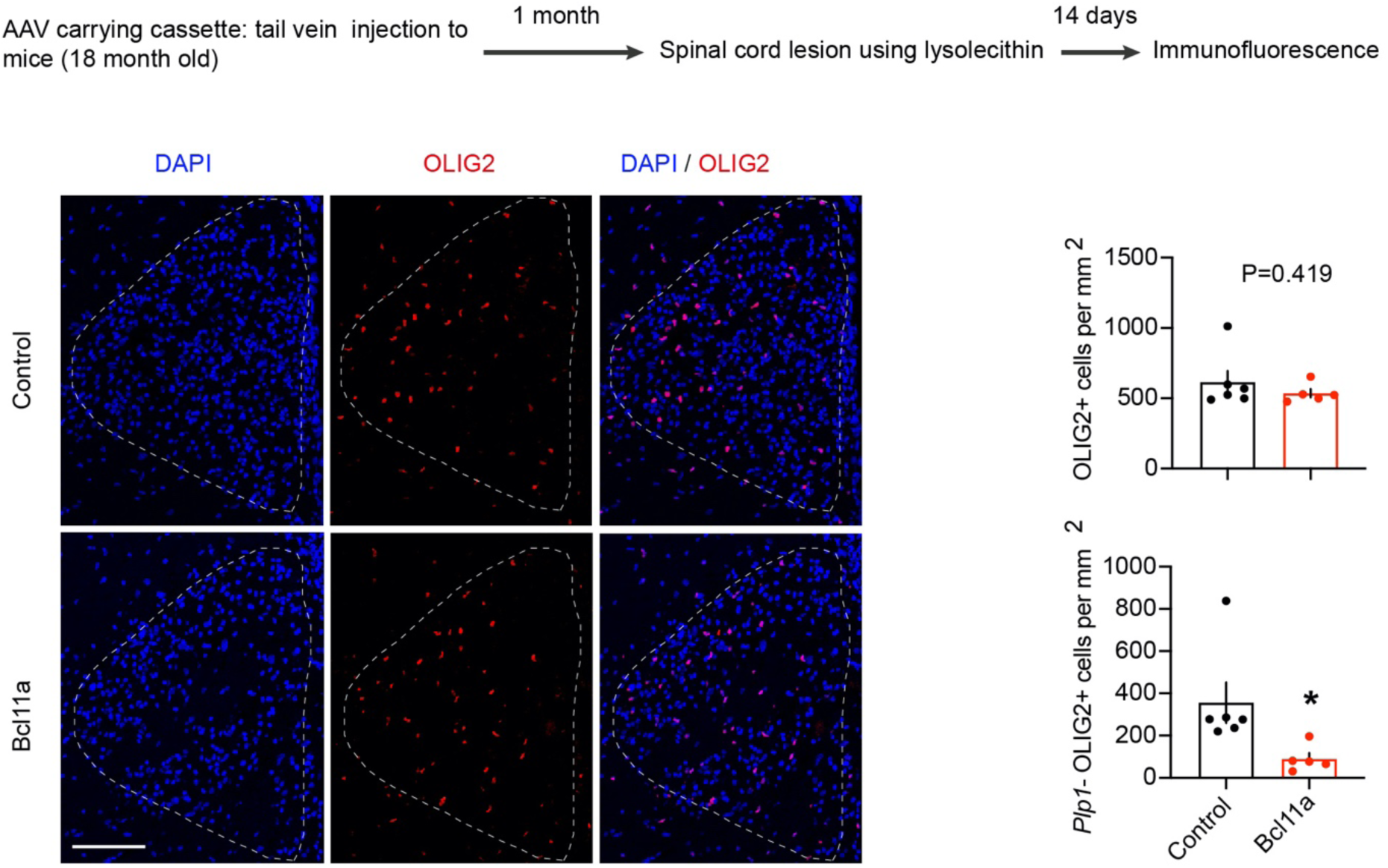
Immunofluorescence analysis using antibody to OLIG2 after OL lineage specific Bcl11a expression in aged mice. The AAV containing the Sox10-driven construct carrying either Gfp alone (control) or Bcl11a linked to Gfp via a self-cleaving peptide element (P2A) was delivered by tail-vein injection (see Figure 6A). In the lesioned area, the number of OLIG2+ cells per mm^2^ was counted and plotted. In the same region, the number of Plp1⁻ OLIG2⁺ cells per mm² was also calculated and plotted (see Figure 6B for *Plp1*⁺ cells per mm²). n=5-6 mice, mean±SEM, P: p-value, ** P<0.01, Student’s t test (unpaired, two tailed). scale bar: 100 µm. The white dotted line demarcates the lesion area.

**Supplementary Table 1**. All switches during OL-lineage progression. (Excel file)

**Supplementary Table 2.** Transcription factor (TF) switches during OL-lineage progression. (Excel file)

**Supplementary Table 3:**
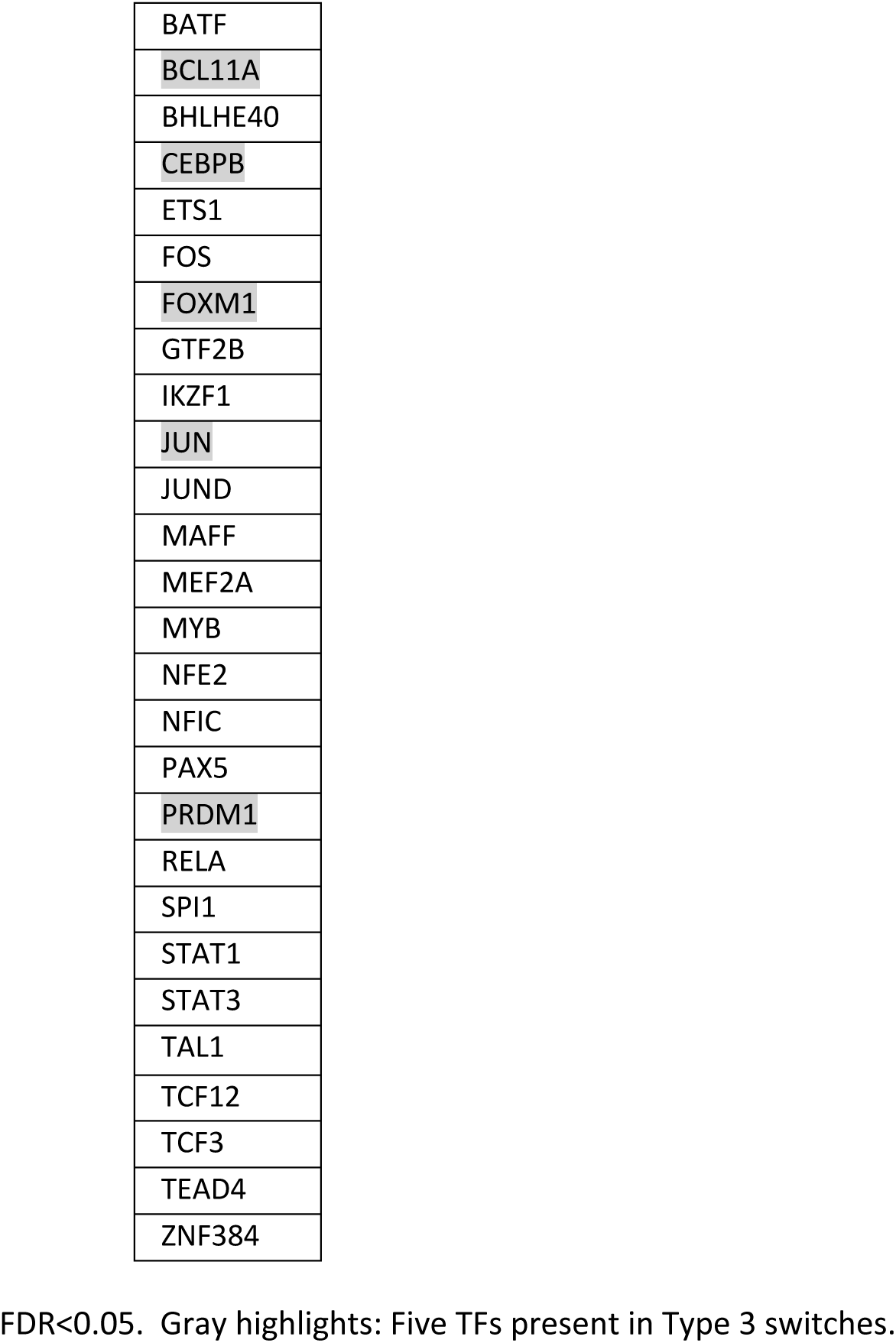
27 Transcription factors (TFs) identified by both Encode and Remap2.

|  |
| --- |
| BATF |
| BCL11A |
| BHLHE40 |
| CEBPB |
| ETS1 |
| FOS |
| FOXM1 |
| GTF2B |
| IKZF1 |
| JUN |
| JUND |
| MAFF |
| MEF2A |
| MYB |
| NFE2 |
| NFIC |
| PAX5 |
| PRDM1 |
| RELA |
| SPI1 |
| STAT1 |
| STAT3 |
| TAL1 |
| TCF12 |
| TCF3 |
| TEAD4 |
| ZNF384 |

FDR<0.05. Gray highlights: Five TFs present in Type 3 switches.

**Supplementary Table 4:** Bcl11a DNA and protein sequence accession numbers for 23 vertebrate species used in this study.

| Species | cds accession | Protein accession |
| --- | --- | --- |
| Human | ENST00000642384.2 | ENSP00000496168 |
| Mouse | ENSMUST00000109514.8 | ENSMUSP00000105140 |
| Rat | ENSRNOT00000106316.1 | ENSRNOP00000093688 |
| Chimpanzee | ENSPTRT00000046502.3 | ENSPTRP00000042480 |
| Gorilla | ENSGGOT00000067107.1 | ENSGGOP00000038984 |
| Elephant | ENSLAFT00000003101.3 | ENSLAFP00000002578 |
| Horse | ENSECAT00000132103.1 | ENSECAP00000085653 |
| Megabat | ENSPVAT00000000041.1 | ENSPVAP00000000040 |
| Macaque | ENSMMUT00000097395.1 | ENSMMUP00000074838 |
| Rabbit | ENSOCUT00000001202.4 | ENSOCUP00000001042 |
| Chicken | ENSGALT00010041405.1 | ENSGALP00010024257 |
| Turkey | ENSMGAT00000014673.3 | ENSMGAP00000013758 |
| Blue-tit | ENSCCET00000017097.1 | ENSCCEP00000010895 |
| Painted turtle | ENSCPBT00000044255.1 | ENSCPBP00000037734 |
| Lizard | ENSPMRT00000007787.1 | ENSPMRP00000007275 |
| *Xenopus tropicalis* | ENSXETT00000097709.2 | ENSXETP00000091923 |
| Zebrafish | ENSDART00000087426.7 | ENSDARP00000081860 |
| Elephant shark | ENSCMIT00000005454.1 | ENSCMIP00000005270 |
| Stickleback | ENSGACT00000017770.1 | ENSGACP00000017735 |
| Atlantic cod | ENSGMOT00000072043.1 | ENSGMOP00000061821 |
| Opossum | ENSMODT00000074969.1 | ENSMODP00000042447 |
| Coelacanth | ENSLACT00000004033.1 | ENSLACP00000003997 |
| Lamprey | XM_032948470.1 | XP_032804361.1 |

cds: coding DNA sequence

## Supplementary Text

Please see **Supplementary Figure 1**. We observed elevated expression of Hes1 in OPCs compared with both PreOLs and OLs, consistent with its role as a Notch effector that maintains the OPC progenitor state and inhibits oligodendrocyte maturation (Brosnan et al, 2009; Ogata et al., 2011).

Compared with PreOLs, adult OPCs isolated from 2–3-month-old rats did not show higher RNA expression of canonical OPC markers: Pdgfra, Pcdh15, and C1ql1. However, as expected, OPCs expressed higher levels of these markers than mature OLs. One possible explanation is the intrinsic heterogeneity of adult OPC populations. Adult OPCs exist in multiple transcriptional states, including quiescent-like and differentiation-primed states. During early differentiation, OPC markers such as Pdgfra may not be immediately extinguished, and PreOLs may transiently retain these transcripts. The PreOL population captured in our study represents intermediate states transitioning from OPCs to OLs, potentially still carrying residual OPC-associated RNAs from activated OPCs. Therefore, comparison of PreOLs with the total heterogeneous OPC pool, which includes quiescent-like OPCs, may give the appearance of higher canonical OPC marker expression in PreOLs.

Among the PreOL-specific markers, Gpr17 clearly distinguished the Pre-OL state in our data, showing higher expression compared with both OPCs and OLs. Bcas1 and Enpp6 showed higher expression in PreOLs compared with OPCs. However, when PreOLs were compared with OLs, Bcas1 appeared lower in PreOLs, whereas Enpp6 expression remained largely unchanged.

The OL markers Mobp and Mog showed significantly higher expression in OLs compared with OPCs, whereas their expression was not altered between OPCs and PreOLs. In contrast, the canonical OL maturity marker Mbp showed a progressive and significant increase during differentiation, with expression levels clearly following the expected pattern: OL > PreOL > OPC.

We did not detect any difference in the astrocyte marker Gfap across these cell-type comparisons, suggesting a similar level of unavoidable astrocytic contamination across groups. Therefore, such contamination is unlikely to confound the interpretation of differential gene expression among OPCs, PreOLs and OLs.

